# Local-Global Parcellation of the Human Cerebral Cortex From Intrinsic Functional Connectivity MRI

**DOI:** 10.1101/135632

**Authors:** Alexander Schaefer, Ru Kong, Evan M. Gordon, Timothy O. Laumann, Xi-Nian Zuo, Avram J. Holmes, Simon B. Eickhoff, B. T. Thomas Yeo

**Author notes:** Address correspondence to: B.T. Thomas Yeo ECE, ASTAR-NUS CIRC, SINAPSE & MNP National University of Singapore.

## Abstract

A central goal in systems neuroscience is the parcellation of the cerebral cortex into discrete neurobiological “atoms”. Resting-state functional magnetic resonance imaging (rs-fMRI) offers the possibility of *in-vivo* human cortical parcellation. Almost all previous parcellations relied on one of two approaches. The local gradient approach detects abrupt transitions in functional connectivity patterns. These transitions potentially reflect cortical areal boundaries defined by histology or visuotopic fMRI. By contrast, the global similarity approach clusters similar functional connectivity patterns regardless of spatial proximity, resulting in parcels with homogeneous (similar) rs-fMRI signals. Here we propose a gradient-weighted Markov Random Field (gwMRF) model integrating local gradient and global similarity approaches. Using task-fMRI and rs-fMRI across diverse acquisition protocols, we found gwMRF parcellations to be more homogeneous than four previously published parcellations. Furthermore, gwMRF parcellations agreed with the boundaries of certain cortical areas defined using histology and visuotopic fMRI. Some parcels captured sub-areal (somatotopic and visuotopic) features that likely reflect distinct computational units within known cortical areas. These results suggest that gwMRF parcellations reveal neurobiologically meaningful features of brain organization and are potentially useful for future applications requiring dimensionality reduction of voxel-wise fMRI data. Multi-resolution parcellations generated from 1489 participants are available (https://github.com/ThomasYeoLab/CBIG/tree/master/stable_projects/brain_parcellation/Schaefer2018_LocalGlobal)

## Introduction

The human cerebral cortex is a thin folded sheet of neural tissue that provides the substrate for sensory and motor functions, and also higher mental processes that distinguish humans from other animals. Complex behavior arises from the transformation of neural signals across networks of distinct cortical areas (Ungerleider and Desimone 1986; Felleman and Van Essen 1991) that are putative atoms of neural processing. Accurate labeling of cortical areas is therefore an important problem in systems neuroscience (Amunts and Zilles 2015).

Cortical areas have traditionally been defined based on the idea that a cortical area should exhibit distinct function, architectonics, connectivity and topography (Kaas 1987; Felleman and Van Essen 1991; Eickhoff and Grefkes 2011). Each of these criteria can be interrogated using a broad range of invasive techniques, including direct electrophysiological recordings during behavioral manipulations to measure function (Shadlen and Newsome, 1998) and topography (Hubel and Wiesel, 1965), histological staining of cell body distribution, myelination and transmitter receptor distribution to measure architectonics (Brodmann 1909; Vogt and Vogt 1919; von Economo and Koskinas 1925; Amunts and Zilles 2015), and retrograde and anterograde tracing via axonal transport to measure neuronal connectivity (Peyron et al. 1997).

Advances in non-invasive brain imaging technologies, such as positron emission topography (PET; Raichle 1987; Petersen et al. 1988) and fMRI (Kwong et al. 1992; Ogawa et al. 1992), offer the possibility of estimating cortical areas *in-vivo* (Sereno et al. 1995; Hinds et al. 2009; Van Essen and Glasser 2014). One promising approach is resting-state functional connectivity (RSFC), which measures synchronization of rs-fMRI signals between brain regions while a subject is lying at rest in the scanner, not performing any directed task (Biswal et al. 1995). Although not a direct measure of anatomical connectivity (Vincent et al. 2007; Honey et al. 2009; Lu et al. 2011), RSFC is sufficiently constrained by anatomy to provide insights into large-scale circuit organization (Fox and Raichle 2007; Buckner et al. 2013) and are strongly associated with task-evoked networks (Smith et al. 2009; Mennes et al. 2010; Cole et al. 2014; Krienen et al. 2014; Bertolero et al. 2015; Yeo et al. 2015a; Tavor et al. 2016). In addition, RSFC is heritable (Glahn et al. 2010; Yang et al. 2016) and correlated with gene expression across the cortical mantle (Hawrylycz et al. 2015; Richiardi et al. 2015; Krienen et al. 2016).

Consequently, rs-fMRI has been widely utilized to estimate large-scale brain networks (Greicius et al. 2003; Beckmann et al. 2005; Vincent et al. 2008; Damoiseaux et al. 2006; De Luca et al. 2006; Fox et al. 2006; Vincent et al. 2006; Dosenbach et al. 2007; Margulies et al. 2007; Seeley et al. 2007; Calhoun et al. 2008; Zuo et al. 2010). While some work has focused on parcellating the brain into a small number of (less than twenty) functional systems (van den Heuvel et al. 2008; Bellec et al. 2010; Power et al. 2011; Lee et al. 2012; Zuo et al. 2012; Hacker et al. 2013), others have focused on finer subdivisions (Cohen et al. 2008; Eickhoff et al. 2011; Craddock et al. 2012; Blumensath et al. 2013; Ryali et al. 2013; Shen et al. 2013; Wig et al. 2014a; Honnorat et al. 2015; Glasser et al. 2016). Given that there is an estimated 300 to 400 classically defined cortical areas in the human cerebral cortex (Van Essen et al. 2012a), these final subdivisions might correspond with more precision to cortical areas (Gordon et al. 2016).

These finer brain parcellations might also capture sub-areal features since classically defined cortical areas are often not homogeneous (Kaas 1987). Well-known intra-areal heterogeneities include occular dominance bands, orientation bands, and cytochrome oxidase dense puffs within primary visual cortex (Livingstone and Hubel 1984). Heterogeneity can be observed even at the macro-scale (i.e., at the resolution of fMRI), such as somatotopy within primary somatosensory cortex (Lotze et al., 2000). Such heterogeneities have prompted a recent proposal to delineate parcels that might be referred to as “neurobiological atoms” (Eickhoff et al., 2017). The difference between traditional cortical areas and neurobiological atoms is especially salient for brain regions that are topographically organized. For example, the different body representations are arguably distinct neurobiological units and might therefore warrant differentiation. Indeed, when modeling a behavioral task with button press, it is probably useful to distinguish between the hand and tongue motor regions. Existing rs-fMRI parcellations (Yeo et al. 2011; Long et al., 2014; Gordon et al. 2016) already appeared to capture certain sub-areal features, including somatotopy and visual eccentricity (Buckner and Yeo 2014).

There are two major approaches to parcellating the brain using rs-fMRI: local gradient and global similarity approaches. The local gradient or boundary mapping approach (Cohen et al. 2008; Nelson et al. 2010; Hirose et al. 2012; Wig et al. 2014b; Laumann et al. 2015; Gordon et al. 2016; Xu et al. 2016) exploits the fact that RSFC patterns can abruptly change from one spatial location to a nearby location. These abrupt changes can be detected by computing local gradients in whole brain RSFC patterns. Previous work suggests that the local gradient approach is especially suited for delineating cortical areas because detecting abrupt changes in RSFC is similar to histological delineation of cortical areas (Cohen et al. 2008; Buckner and Yeo 2014; Wig et al. 2014b).

In contrast, the global similarity approach (e.g., mixture models and spectral clustering) cluster brain regions based on similarity in rs-fMRI time courses or RSFC patterns (van den Heuvel et al. 2008; Power et al. 2011; Yeo et al. 2011; Craddock et al. 2012; Ryali et al. 2013; Shen et al. 2013; Honnorat et al. 2015). The global similarity approaches often disregard spatial distance (e.g., medial prefrontal and posterior cingulate regions can be grouped into the same default network despite their spatial distance), although some approaches discourage spatially disconnected parcels (Honnorat et al. 2015) or neighboring brain locations to have different parcellation labels (Ryali et al. 2013). Because the global similarity approach seeks to group voxels or vertices with similar rs-fMRI time courses or RSFC patterns, the resulting parcels might be highly connectionally homogeneous, suggesting that the parcels might reflect putative neurobiological atoms (Eickhoff et al., 2017). Just as importantly, the parcellations might potentially be utilized as a dimensionality reduction tool in many rs-fMRI applications, where dealing with the fMRI data at the original voxel level resolution is difficult (Finn et al. 2015; Smith et al. 2015). In these applications, each parcel could be represented by an average time course of voxels or vertices within the parcel. Therefore the voxels or vertices constituting a parcel should ideally have very similar fMRI time courses.

Local gradient approaches do generate homogeneous parcels (Gordon et al. 2016) because they implicitly encourage homogeneity by discouraging high RSFC gradients within a parcel. However, because global similarity approaches explicitly optimize for connectional homogeneity, they should in theory be able to produce more homogeneous parcels than local gradient approaches. On the other hand, local gradient approaches appear to be more sensitive to certain biological boundaries than global similarity approaches. For example, boundary between areas 3 and 4 is captured by local gradient approaches (Gordon et al. 2016), but not global similarity approaches (Craddock et al. 2012; Shen et al. 2013). Therefore integrating local gradient and global similarity approaches might generate parcellations that are both neurobiological meaningful and useful for future applications requiring dimensionality reduction.

In this work, we developed a gradient-weighted Markov Random Field (gwMRF) model that fuses the local gradient and global similarity approaches. MRFs are used in many neuroimaging software packages for delineating anatomical structures (Zhang et al., 2001; Fischl et al., 2002). For example, the FSL FAST (Zhang et al., 2001) MRF model balances a global similarity objective that encourages voxels with similar T1 intensities to be labeled the same structure with a local smoothness objective that encourages neighboring voxels to have the same labels (Zhang et al., 2001). The key idea in this paper is to modify the local smoothness objective to encourage neighboring voxels to have the same labels only in the presence of low local RSFC gradients. The resulting gwMRF fusion model was compared with four publicly available parcellations (Craddock et al. 2012; Shen et al. 2013; Glasser et al. 2016; Gordon et al. 2016) using multiple multimodal datasets across MNI, fsaverage and fsLR coordinate systems. After controlling for the number of parcels, our results suggest that compared with other fully automatic approaches, parcellations generated by the gwMRF approach enjoy greater functional and connectional homogeneity as measured by task-fMRI and rs-fMRI respectively, while achieving comparable agreement with architectonic and visuotopic boundaries. Finally, we applied the gwMRF model to 1,489 participants from the Genomics Superstruct Project (GSP; Holmes et al., 2015) yielding 400, 600, 800 and 1000 parcels, which are publicly available as reference maps for future studies (https://github.com/ThomasYeoLab/CBIG/tree/master/stable_projects/brain_parcellation/Schaefer2018_LocalGlobal).

## Methods

### Overview

A gwMRF parcellation procedure was developed and applied to a rs-fMRI dataset (N = 744) from the Genomics Superstruct Project. The resulting parcellations were compared with four previously published rs-fMRI parcellations using multimodal data from multiple scanners with diverse acquisition and processing protocols. A final set of gwMRF parcellations at different spatial resolutions were estimated from the full GSP dataset (N = 1489) and further characterized.

### Genomics Superstruct Project (GSP) Data

The primary dataset utilized in this work consisted of structural MRI and rs-fMRI data from 1,489 young adults (ages 18 to 35) of the Genomics Superstruct Project (GSP; Holmes et al., 2015). All imaging data were collected on matched 3T Tim Trio scanners (Siemens Healthcare, Erlangen, Germany) at Harvard University and Massachusetts General Hospital using the vendor-supplied 12-channel phased-array head coil. One or two fMRI runs were acquired per participant – 1,083 participants had two runs, 406 participants had one run. Each run was acquired in 3mm isotropic resolution with a TR of 3.0 seconds and lasted for 6 minutes and 12 seconds. The structural data consisted of one 1.2mm isotropic scan for each subject.

Details of the data collection and preprocessing can be found elsewhere (Yeo et al. 2011; Holmes et al. 2015). Briefly, structural scans were processed using FreeSurfer 4.5.0 (http://surfer.nmr.mgh.harvard.edu; Fischl 2012) and structure-function registration was performed using FsFast (http://surfer.nmr.mgh.harvard.edu/fswiki/FsFast; Greve and Fischl 2009). Functional preprocessing steps included slice time correction, motion correction, low pass temporal filtering retaining frequencies below 0.08Hz, regression of motion parameters, ventricular signal, white matter signal, whole brain signal, and linear trend. The final data was projected onto the fsaverage6 surface space (where vertex spacing is roughly 2mm) and smoothed using a 6mm full-width half-maximum kernel. Preprocessing of the GSP data followed the official data release publication (Holmes et al. 2015). However, we recognize that certain preprocessing steps (e.g., whole brain signal regression) are controversial. Therefore additional rs-fMRI datasets preprocessed with alternate pipelines were employed to evaluate the parcellations (see *Parcellation Evaluation Metrics* section). The full GSP dataset was divided into training (N = 744) and test (N = 745) sets.

### Gradient-weighted Markov Random Field (gwMRF) Parcellation

MRF models (Geman and Geman, 1984) are utilized in many well-known neuroimaging software packages for segmenting anatomical brain regions, including FreeSurfer (Fischl et al. 2002; Fischl et al., 2004) and FSL (Zhang et al., 2001). In these software packages, the typical goal is to assign an anatomical label (e.g., white matter, gray matter, cerebrospinal fluid and background) to all brain locations.

A typical MRF model consists of an objective function with multiple competing terms, encoding tradeoffs between ideal properties of the final segmentation (Fischl et al., 2002; Fischl et al., 2004). The objective function almost always contains a global similarity term that encourages brain locations with similar image intensities to be assigned a shared label regardless of spatial proximity (Zhang et al., 2001). For instance, all white matter voxels should have high signal intensities in T1 MRI. If the objective function consists of only the global similarity term, then the MRF is also known as a mixture model, which is a superclass of the particular global similarity approach employed in Yeo et al. (2011).

In addition to the global similarity term, MRF objective functions almost always include a second “pairwise” term that encodes desired relationships between adjacent brain locations. As an example, a common pairwise term (known as the Potts model) encourages neighboring brain locations to have the same segmentation label (Zhang et al., 2001; Ryali et al., 2013). This pairwise term is useful for handling noise inherent in MRI. For example, a white matter voxel might exhibit abnormally low intensity, resulting in an incorrect label if the objective function consists of only the global similarity term. However, in cases where the voxel is surrounded by correctly labeled white matter voxels, the additional pairwise Potts model might overcome the global similarity term, and generate an accurate label assignment.

Our gwMRF parcellation procedure utilizes three terms in the MRF objective function. The first term in the objective function is the analogue of the von Mises-Fisher mixture model employed in Yeo et al. (2011), and thus encodes the global similarity approach. When applied to preprocessed fMRI data from a group of participants, the fMRI time courses are concatenated after normalization to mean of zero and standard deviation of one. The resulting global similarity term encourages brain locations with similar fMRI time courses to be assigned to the same parcel. Our particular choice of time course similarity is equivalent to the Pearson product-moment correlation coefficient computed for each participant and then averaged across participants (see Supplementary Methods S1 for details).

The second term in the MRF objective function is a pairwise term that encodes the local gradient approach. Unlike the Potts model, which penalizes any adjacent pairs of brain locations with different parcellation labels, our pairwise term down weighs the penalties in the presence of strong local RSFC gradients (hence the name “gradient weighted”). In this work, we utilize state-of-the-art RSFC gradients computed by Gordon et al. (2016), where the gradients ranged in values from zero to one. As the gradient magnitude between two adjacent brain locations increases from zero to one, the penalty of the two brain locations having different parcellation labels decreases exponentially to zero.

If the MRF objective function contains only the previous two terms, then the resulting parcellation will contain many spatially distributed parcels because of strong long-range RSFC. Requiring parcels to be spatially connected in a MRF framework (with minimal other assumptions) is non-trivial (Nowozin and Lampert 2010; Honnorat et al. 2015). Here we include a third spatial connectedness term in the MRF objective function, which encourages brain locations constituting a parcel to not be too far from the parcel center. If the third term in our MRF objective function is sufficiently strong, then parcels will indeed become spatially connected. This approach is significantly less computationally expensive than competing approaches (Nowozin and Lampert 2010; Honnorat et al. 2015).

More details of the gwMRF model (objective function) can be found in Supplementary Methods S1. Given the MRF objective function and data from a cortical hemisphere, a final parcellation can be obtained by optimizing the objective function with various standard techniques. Here we utilize graph cuts (Delong et al., 2010) within a maximum-a-posteriori (MAP) estimation framework. Readers should note that the third term in the objective function exists to ensure parcels are spatially connected. However, an overly strong third term would result in extremely round parcels, which are not biologically plausible. For example, we expect cortical areas in the cingulate to be long and narrow (Vogt 2009). To avoid this issue, the optimization procedure starts with a strong weight on the third term and then gradually decreases the weight of this third term. While there is no theoretical guarantee, we find that the third term is driven to zero for almost all parcels in practice. Details of our optimization procedure can be found in Supplementary Methods S2. The parcellation code is publicly available (https://github.com/ThomasYeoLab/CBIG/tree/master/stable_projects/brain_parcellation/Scha_efer2018_LocalGlobal)

### Parcellation Evaluation Metrics

If the estimated parcels were indeed neurobiological atoms of the cerebral cortex, then each parcel should have uniform (homogeneous) function and connectivity, as well as correspond well with boundaries of cortical areas delineated with architectonics and topography. Therefore to assess the quality of a given cerebral cortex parcellation, evaluation metrics based on multimodal measurements of function, architectonics, connectivity and visuotopy were considered. The evaluation metrics will be utilized to compare gwMRF parcellations with other publicly available parcellations after controlling for the number of parcels (next section).

1. *Architectonics.* Ten human architectonic areas 17, 18, 1, 2, 3 (combining areas 3a and 3b), 4 (combining areas 4a and 4p), 6, hOc5, 44 and 45 (Geyer et al. 1996, 1999, 2000, 2001; Amunts et al. 1999, 2000, 2004; Geyer 2004; Malikovic et al. 2007) were considered. The histological areas were mapped to fsLR surface space by Van Essen et al. (2012a) based on Fischl et al. (2008). Alignment between the architectonic and parcellation boundaries was numerically assessed using average Hausdorff distance (Yeo et al. 2010). Briefly, for each boundary vertex of each architectonic area, the geodesic distance to the closest parcellation boundary was computed. The geodesic distances were averaged across all boundary vertices of an architectonic area, resulting in a Hausdorff distance for each architectonic area (shown in Figure 2 of *Results).* Smaller Hausdorff distance indicates better agreement between parcellation and histological boundaries. When comparing between parcellations, paired-sample t-test (dof = 19) was performed on the Hausdorff distances on both hemispheres.
2. *Visuotopy.* 18 retinotopic areas in fsLR space from a previous study were considered (Abdollahi et al. 2014). The retinotopic areas were obtained with fMRI visuotopic mapping in individual subjects and then transformed and averaged in fsLR space using multimodal surface matching (Abdollahi et al. 2014). One tricky issue is that stimulating peripheral visual field is hard, and hence visuotopic maps tend to span about half the true extent of each area (Hinds et al. 2009). Consequently, only boundary vertices bordering adjacent retinotopic areas were considered as valid boundary vertices. In total, there were 46 pairs of adjacent retinotopic areas for both hemispheres. Agreement between the visutopic and parcellation boundaries was assessed using average Hausdorff distance. For each valid visuotopic boundary vertex, the geodesic distance to the closest parcellation boundary vertex was computed. The geodesic distances were averaged across all boundary vertices of each adjacent pair of retinotopic areas (e.g., V1 and V2), and then averaged across all 46 pairs of retinotopic areas, resulting in a single average distance for all retinotopic areas shown in Figure 5 of *Results.* When comparing parcellations, paired-sample t-test (dof = 45) was performed on the hausdorff distances.
3. *Function.* Task-fMRI data from the Human Connectome Project (HCP) S900 release in fsLR surface space (Barch et al. 2013) was utilized to evaluate the functional homogeneity of the parcels. This dataset comprises seven cognitive domains: social cognition, motor, gambling, working memory, language processing, emotional processing and relational processing. The subset of subjects (N = 745) with available z-maps for all contrasts was considered. To assess the functional inhomogeneity of a parcellation, the standard deviation of z-values for each parcel was computed for each task contrast. A lower standard deviation indicates higher functional homogeneity within the parcel. The standard deviations were averaged over all parcels while accounting for parcel size:

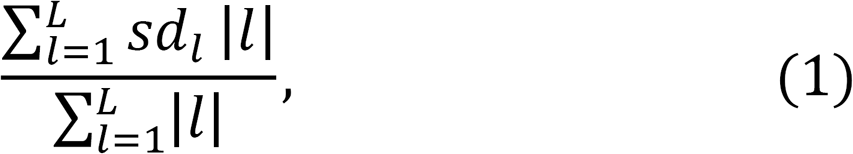

where *sd_l_* is the standard deviation of task activation z-values for parcel *l* and |*l*| is the number of vertices in parcel *l*. The functional inhomogeneity metric ((Eq. (1)) was computed separately for each participant and each task contrast and then averaged across all contrasts within a cognitive domain, resulting in a functional inhomogeneity measure per cognitive domain (shown in Figure 3 of *Results).* The number of task contrasts per cognitive domain ranges from three for the emotion domain to eight for the working memory domain. When comparing between parcellations, the inhomogeneity metric (Eq. (1)) was averaged across all contrasts within a cognitive domain and then averaged across all seven cognitive domains before a paired-sample t-test (dof = 744) was performed.
4. *Connectivity.* To assess the functional connectivity homogeneity of a parcellation, we utilized rs-fMRI data from three different datasets. The first dataset is the GSP test set (N = 745) discussed in the *Genomics Superstruct Project (GSP) Data* section. To assess whether a parcellation would generalize well to data from different scanners, acquisition protocols, preprocessing strategies, coordinate systems and population groups, we considered two additional datasets. The first additional dataset was the group-averaged connectivity matrix (N = 820) in fsLR space from the HCP S900 release (Van Essen et al. 2012b; Smith et al. 2013). Details of the HCP data collection, preprocessing and functional connectivity matrix computation can be found elsewhere (HCP S900 manual; Van Essen et al. 2012b; Glasser et al. 2013; Smith et al. 2013). The main differences between the HCP and GSP data were that the HCP data were collected on a custom-made Siemens 3T Skyra scanner using a multiband sequence, and denoised using ICA-FIX (Griffanti et al. 2014) and Wishart soft thresholding with minimal (2mm) smoothing (HCP S900 manual; Glasser et al. 2016).

**Figure 2.**
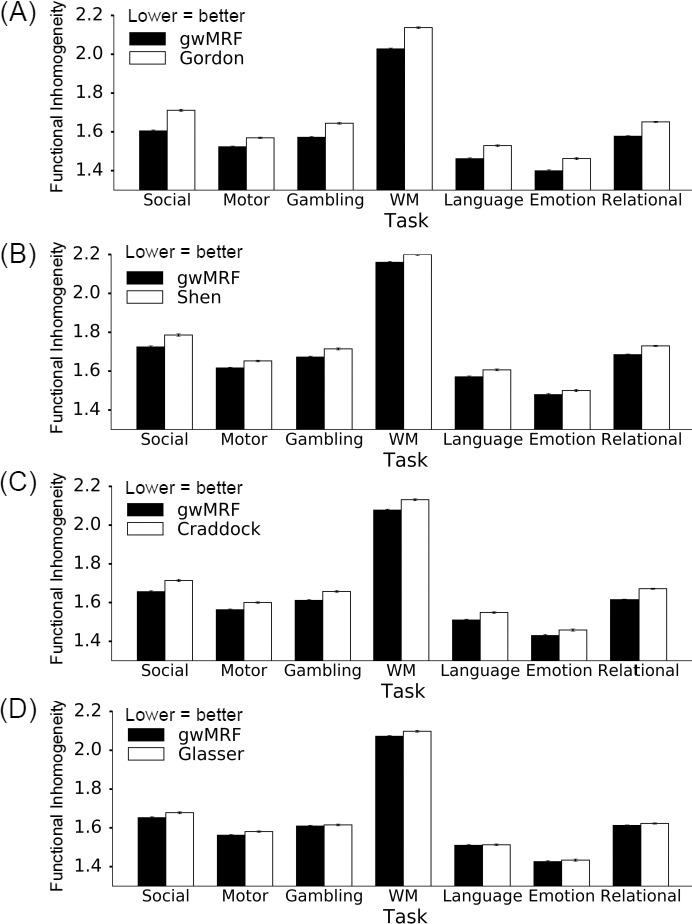
Functional inhomogeneity measured by standard deviation of fMRI task activation within each parcel, and averaged across all parcels and contrasts of each cognitive domain (N=745). Lower standard deviation indicates higher functional homogeneity. Our approach generated parcellations more functionally homogeneous than (A) Gordon (p ≈ 0), (B) Shen (p ≈ 0), (C) Craddock (p ≈ 0) and (D) Glasser (p = 4.8e-229). It is worth mentioning that the Glasser parcellation was partially derived from this dataset.

**Figure 3.**
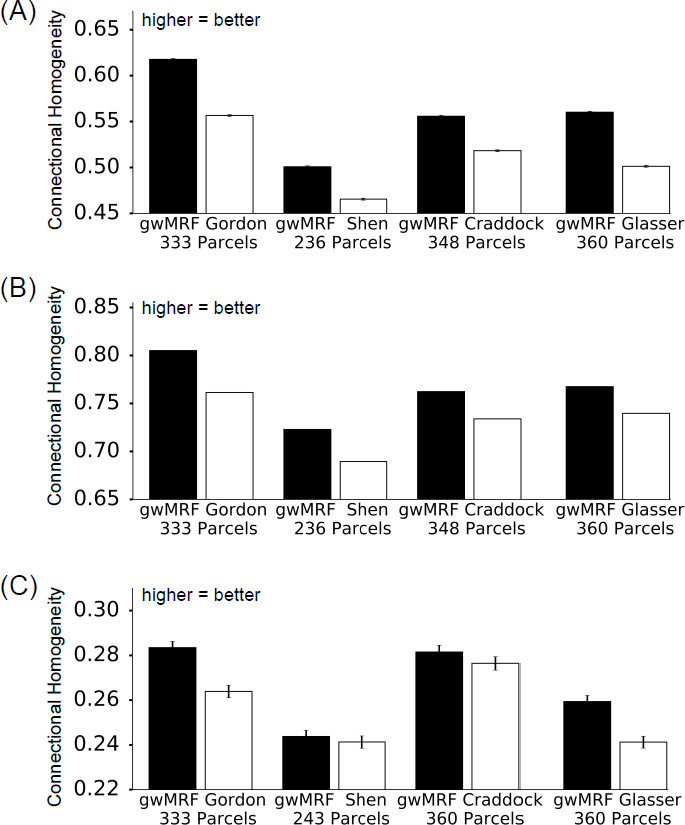
Connectional homogeneity as measured by rs-fMRI computed on the (A) GSP test set (N = 745) in fsaverage space, (B) HCP dataset (N = 820) in FSLR space, and (C) NKI dataset (N = 205) in MNI space. The gwMRF fusion approach generated parcellations that achieved better connectional homogeneity than all other parcellations across all three datasets (p < 2.8e-28 for all comparisons). There is no error bar for the HCP data because the HCP group-average dense connectivity matrix was utilized. It is worth mentioning that the Glasser parcellation was partially
derived from the HCP dataset.

**Figure 5.**
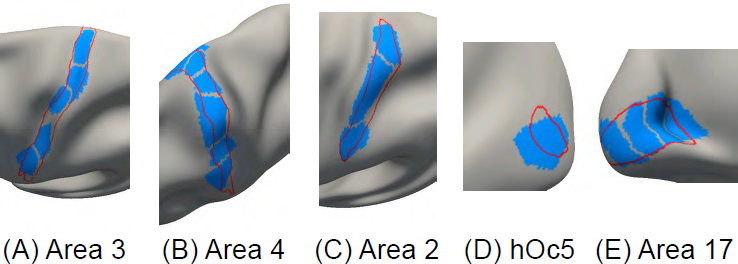
Parcels (blue) of the 400-area cerebral cortex parcellation overlaid on histological (red) boundaries of left hemisphere (A) area 3, (B) area 4, (C) area 2, (D) hOc5 and (E) area 17. Other histological areas are found in Figure S9.

The second dataset consisted of 205 subjects from the Nathan Kline Institute (NKI) Rockland Sample Release 1-3 (Nooner et al., 2012) normalized to MNI volumetric space. The NKI dataset have been employed in several recent lifespan connectomics studies (Betzel et al. 2014; Cao et al. 2014; Yang et al. 2014; Jiang et al. 2015; Xu et al. 2015). The major difference between the GSP and NKI datasets was that the NKI dataset consisted of participants ranging from ages 6 to 85, offering the opportunity to assess the performance of the parcellations across the human lifespan (Zuo et al., 2017). Two major differences between the GSP and NKI preprocessing were that anatomical CompCor (instead of whole brain signal regression) and no smoothing were applied to the NKI dataset (Behzadi et al. 2007; Schaefer et al. 2014). Details of the NKI data collection and preprocessing can be found elsewhere (Nooner et al. 2012; Schaefer et al. 2014). The preprocessing pipeline can be downloaded from https://github.com/alexschaefer83/DynamicHubs/

Functional connectivity homogeneity was computed by averaging Pearson moment-product correlation coefficients between all pairs of vertices (or voxels) within each parcel. The average correlations are then averaged across all parcels while accounting for parcel size:

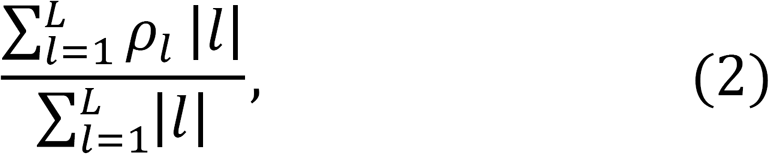

where *ρ_l_* is the functional connectivity homogeneity of parcel *l* and |*l*| is the number of vertices (or voxels) for parcel *l*.

The functional connectivity homogeneity metric (Eq. (2)) was computed for each participant separately and averaged across participants, except for the HCP group-averaged connectivity matrix where inter-subject averaging has already been performed. The resulting functional connectivity homogeneity measures are shown in Figure 4 of *Results.* When comparing between parcellations, the homogeneity metric (Eq. (2)) was computed for each participant separately before a paired-sample t-test (dof = 744 for GSP, dof = 204 for NKI) was performed. Statistical test could not be performed for the HCP data because the HCP group-averaged connectivity matrix was utilized.

**Figure 4.**
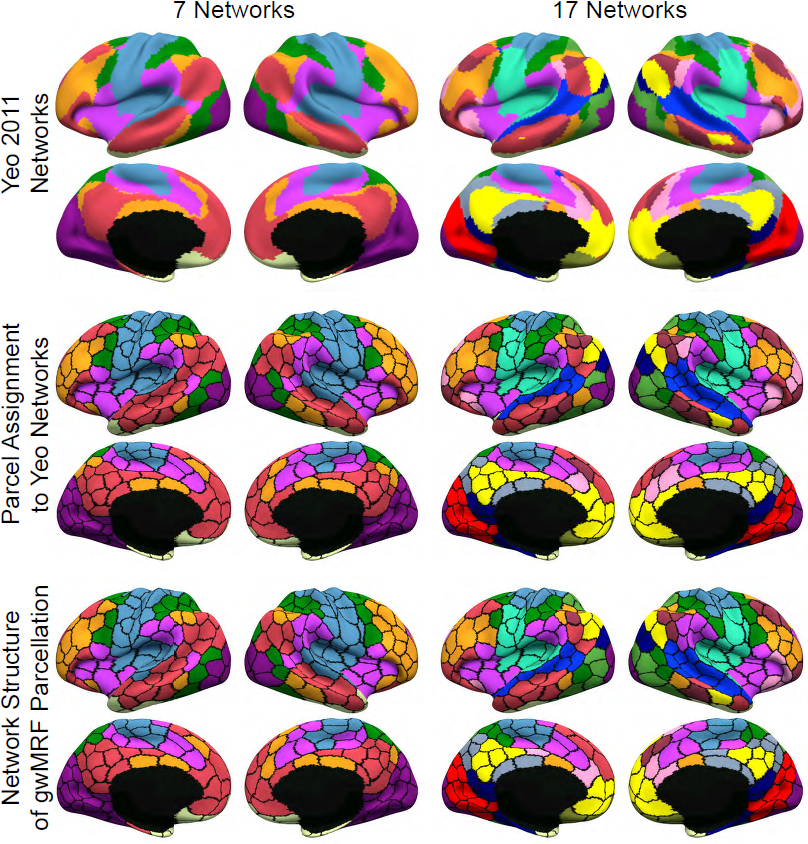
Network structure is preserved in the 400-area parcellation. First row shows 7 and 17 networks from Yeo et al. (2011). Second row shows each gwMRF parcel assigned a network color based on spatial overlap with networks from Yeo et al. (2011). Last row shows community structure of gwMRF parcellation after clustering. Observe striking similarity between second and third row, suggesting that network structure of the original resolution data is preserved in the 400-area parcellation. Results for 600-area, 800-area and 1000-area parcellations can be found in Figures S6, S7 and S8.

We note that a cortical parcellation with more parcels will on average perform better on the proposed evaluation metrics. The reason is that a cortical parcellation with more parcels will have smaller parcels on average. Smaller parcels are likely to be more functionally and connectionally homogeneous. Similarly, more parcels lead to more boundary vertices, and would therefore enjoy better (lower) architectonic and visuotopic Hausdorff distance (on average. Therefore, when comparing parcellations, it is important to control for the number of parcels as discussed in the next section.

### Comparison with Other Parcellations

To evaluate the quality of our parcellation procedure, the gwMRF model was applied to the GSP training set (N = 744). The setting of various “free” parameters in the gwMRF model can be found in Supplementary Methods S6. The resulting gwMRF parcellations were compared to four publicly available parcellations (Craddock et al. 2012; Shen et al. 2013; Glasser et al. 2016; Gordon et al. 2016) using criteria described in the *Parcellation Evaluation Metrics* section. Three of the parcellations were generated using fully automatic algorithms from rs-fMRI (Craddock et al. 2012; Shen et al. 2013; Gordon et al. 2016), while one parcellation was generated from the application of a semi-automatic algorithm to multimodal data, including rs-fMRI, relative myelin mapping and task-fMRI (Glasser et al., 2016). The semi-automatic algorithm required an anatomist to manually select multimodal gradients that fitted prior anatomical knowledge of cortical areas. The Gordon and Glasser parcellations consisted of 333 and 360 parcels respectively. Parcellations of multiple resolutions were available for Craddock et al. (2012) and Shen et al. (2013). Since it has been estimated that there are between 300 to 400 human cortical areas (Van Essen et al. 2012a), the Shen and Craddock parcellations with 300 and 400 parcels respectively were considered.

There were several issues worth elaborating when comparing brain parcellations. The first issue was the number of parcels *L.* As discussed in the previous section, as the number of parcels increases, the parcellation will on average perform better in all the proposed evaluation metrics. To control for the number of parcels *L,* the gwMRF parcellation procedure was run with different number of parcels to match the four publicly available parcellations. In the case of the Gordon parcellation, there were vertices between parcels not assigned to any parcellation label, which generally increased the performance of the parcellation on all evaluation metrics. Therefore, boundary vertices with the worst connectional homogeneity in the GSP training set were removed from the gwMRF parcellation to match the number of unlabeled vertices in the Gordon parcellation.

The second important issue is that the parcellations were in different coordinate systems. Gordon and Glasser parcellations were in fsLR surface space, Shen and Craddock parcellations were in MNI volumetric space, and the gwMRF parcellations were in fsaverage space. Similarly, the evaluation data were in different coordinate systems. The architectonic, visuotopic and task fMRI data were in fsLR surface space. The GSP rs-fMRI data were in fsaverage space, the HCP rs-fMRI data were in fsLR surface space, while the NKI rs-fMRI data were in MNI152 volumetric space. For evaluation, the different parcellations were transformed to the coordinate system where the evaluation data resided. For example, to evaluate connectional homogeneity using the GSP test set (which resided in fsaverage surface space), the Gordon, Glasser, Shen and Craddock parcellations were transformed to fsaverage surface space. As another example, to evaluate alignment with architectonic and visuotopic boundaries (which resided in fsLR surface space), the Shen, Craddock and gwMRF parcellations were transformed into fsLR space. The nonlinear transformations between fsLR and fsaverage spaces, as well as between fsaverage and MNI spaces are detailed elsewhere (Buckner et al. 2011; Van Essen et al., 2012a; http://sumsdb.wustl.edu/sums/directory.do?id=8291757&dir_name=Inter-atlas_deformation_maps).

The two issues (number of parcels and coordinate systems) actually interacted because transforming a parcellation to a target coordinate system might affect the resulting number of parcels. For example, the Shen and Craddock parcellations included subcortical structures. Mapping their parcellations to fsaverage (or fsLR) space reduced the number of parcels, resulting in 236 and 348 parcels for the Shen and Craddock parcellations respectively. For comparisons in fsaverage (or fsLR) space, the gwMRF parcellation algorithm was run to ensure the same number of parcels as the Shen and Craddock parcellations. On the other hand, when comparing the gwMRF parcellations to the Shen or Craddock parcellation in MNI space, a masking procedure ensured the parcellations covered the same aspects of the cerebral cortex. This masking procedure modified the number of parcels, resulting in 243 and 360 parcels for the Shen and Craddock parcellations respectively. For comparisons in MNI space, the gwMRF parcellation algorithm was run to ensure the same number of parcels as the Shen and Craddock parcellations.

Transforming the Shen and Craddock parcellations to the surface spaces was technically challenging. Despite the highly accurate mapping between MNI and fsaverage space developed in Buckner et al. (2011), the transformed parcellations in surface space tended to have rough borders resulting in reduced functional homogeneity. To alleviate such biases, the parcellation boundaries were smoothed on the surface space. The post-processing procedure served to improve the performance of the transformed parcellations and the resulting parcellations appeared visually appealing (Figure S1). Nevertheless, all biases could not be fully removed and so a parcellation created and evaluated in coordinate system X would have inherent advantages in its native space over parcellations created in a different coordinate system and then projected to coordinate system X. Therefore the results should be interpreted with care.

### Stability of gwMRF Parcellations with 400, 600, 800 and 1000 Areas

The gwMRF parcellation algorithm was applied to the GSP training and test sets independently. Parcellations with 400, 600, 800 and 1000 areas were obtained. The setting of various “free” parameters in the gwMRF model can be found in Supplementary Methods S6. To visually appreciate agreement between training and test parcellations, boundaries of both parcellations were overlaid on fsLR inflated surfaces. Quantitative agreement between the training and test parcellations was assessed based on the percentage of vertices assigned to the same parcels.

### Cerebral Cortex Parcellation of 1,489 participants

The gwMRF parcellation algorithm was applied to the full GSP dataset (N = 1,489) to generate a population atlas of cerebral cortical parcellation, which we have made publicly available (https://github.com/ThomasYeoLab/CBIG/tree/master/stable_projects/brain_parcellation/Scha_efer2018_LocalGlobal). Given the hierarchical organization of the cerebral cortex, we do not believe that a single-resolution parcellation will be optimal across all applications. Consequently, we generated parcellations with 400, 600, 800 and 1000 areas. The parcellations are available in fsaverage and fsLR surface spaces, as well as MNI152 volumetric space.

### Network structure of gwMRF parcellations

To investigate whether the parcellations preserved the community structure of the original data, the parcellations were clustered into 7 and 17 networks using a similar procedure to Yeo et al. (2011) applied to the full GSP dataset. For a *L*-area parcellation (where *L* is 400, 600, 800 or 1000), an average time course for each parcel and each subject was computed. For each subject, an *L* × *L* correlation matrix was then computed using the averaged time courses. Similar to Yeo et al. (2011), each correlation matrix was binarized (by keeping only the top 10% of the correlation entries) and averaged across the 1489 subjects. The resulting binarized and averaged correlation matrix was clustered into 7 and 17 networks using the von Mises-Fisher mixture model and compared with the networks from Yeo et al. (2011).

### Further characterization of the 400-area parcellation

The 400-area parcellation was further characterized by juxtaposing the parcels against boundaries of architectonic and visuotopic areas, as well as visualizing its relationship with task activation and connectional homogeneity. We also compared the distribution of parcel volumes against the four previously published parcellations.

## Results

### Local gradient and global similarity approaches capture different aspects of architectonic boundaries

Figure 1 overlays the boundaries of histologically defined areas 3 and 44 (Amunts et al. 1999; Geyer et al. 1999; Fischl et al. 2008; Van Essen et al. 2012a) on the parcels generated from three different approaches. The local gradient approach corresponded to the 333-area Gordon parcellation (Gordon et al., 2016). The gwMRF fusion approach corresponded to the application of the gwMRF parcellation procedure to the GSP training set with 333 parcels (see *Methods).* The global similarity approach corresponded to the gwMRF algorithm with the pairwise term in the objective function removed (i.e., the local gradients have no influence on the resulting parcellation), with all other parameters the same as the gwMRF fusion parcellation.

**Figure 1.**
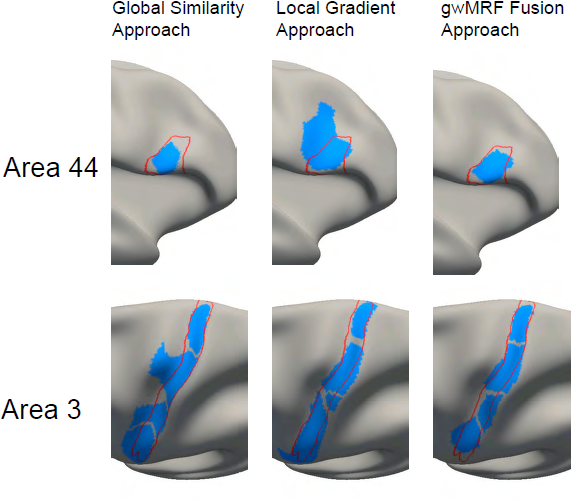
Parcels (blue) from 333-area parcellations using global similarity, local gradient and gwMRF approaches overlaid on histological (red) boundaries of areas 44 and 3. The local gradient approach overestimates the posterior boundary of area 44. On the other hand, the ″bleeding″ of cortical parcels across the central sulcus (area 3) could not be easily avoided without taking into account local gradients. Parcels from the gwMRF fusion approach agree well with both areas.

Compared with the global similarity approach, parcels generated by the local gradient approach agreed better with area 3 boundary (bottom row of Figure 1). Geodesic (average Hausdorff) distance was 4.7mm for the local gradient approach compared with 5.1mm for the global similarity approach. Lower distance indicates better agreement. The “bleeding” of cortical parcels across the central sulcus could not be easily avoided without taking into account the local gradients. On the other hand, parcels generated by the local gradient approach overestimated the posterior boundary of area 44, while the global similarity approach underestimated the posterior and anterior portions of area 44 (top row of Figure 2). Geodesic distance was 5.8mm for the local gradient approach and 6.6mm for the global similarity approach. By fusing the local and global approaches, the resulting parcellation agreed well with both areas 3 and 44 with geodesic distances of 3.9mm and 4.2mm respectively.

### Compared with other fully automatic approaches, the gwMRF model yielded parcellations with comparable architectonic alignment

Figure S1 shows four publicly available parcellations (Craddock, Shen, Gordon and Glasser) and corresponding gwMRF parcellations with matching number of parcels. Figure S2 shows the geodesic distances between the boundaries of ten histologically defined architectonic areas (Geyer et al. 1996, 1999, 2000, 2001; Amunts et al. 1999, 2000, 2004; Geyer 2004; Malikovic et al. 2007) and the boundaries of parcellations generated by the gwMRF model and four publicly available parcellations. Lower distance indicates better agreement (details in *Methods*).

Across ten histological areas, the gwMRF parcellations achieved comparable architectonic distance with Gordon (p = 0.076 uncorrected), better distance than Craddock (p = 0.049 uncorrected), better distance than Shen (p = 2.7e-5 uncorrected) and worse distance than Glasser (p = 0.042 uncorrected).

Relative to the gwMRF parcellation, the Glasser parcellation performed especially well for certain early sensory and late motor areas (areas 17, 18, hOc5, 3, 4). Relative to the Glasser parcellation, the gwMRF parcellation performed especially well for early somatosensory area 2 and was slightly better in putative language areas 44 and 45. The Glasser parcellation utilized a semi-automatic approach requiring an anatomist to manually select specific multimodal information that matches prior knowledge of areal boundaries. The variable differences across architectonic areas might reflect the priorities of the anatomist.

### Compared with other fully automatic approaches, the gwMRF model yielded parcellations with comparable visuotopic alignment

Figure S3 shows the geodesic distances between the boundaries of 18 visuotopically-mapped retinotopic areas from a previous study (Abdollahi et al. 2014) and the boundaries of parcellations generated by the gwMRF model and four publicly available parcellations. Lower distance indicates better agreement (details in *Methods*).

Across 18 visuotopic areas, the gwMRF parcellations achieved lower visuotopic distance than Gordon (p = 8.7e-3 uncorrected), comparable distance with Craddock (p = 0.37 uncorrected), comparable distance with Shen (p = 0.21 uncorrected) and worse distance than Glasser (p = 1.8e-6 uncorrected). As noted above, the Glasser parcellation utilized an approach that semi-automatically selected specific multimodal information to match prior knowledge of areal boundaries.

### Compared with other approaches, the gwMRF model yielded parcellations with higher functional homogeneity

Figure 2 compares the functional inhomogeneity of parcellations generated by the gwMRF model with four publicly available parcellations. Using the HCP task-fMRI data (N = 745), functional inhomogeneity was measured by computing standard deviation of fMRI task activation (z-scores) within each parcel, and then averaged across all parcels and contrasts of a cognitive domain (details in *Methods*). Comparisons between the subplots (e.g., Figures 2A and 2B) are not meaningful because the number of parcels is different across the parcellations.

The gwMRF model generated parcellations that were significantly more functionally homogeneous than Gordon (p ~ 0), Craddock (p ~ 0), Shen (p ~ 0) and Glasser (p = 4.8e-229) with average improvements of 4.7% (Gordon), 2.7% (Craddock), 2.3% (Shen), and 0.8% (Glasser). It is worth mentioning that the Glasser parcellation enjoyed inherent advantage in this evaluation because the parcellation was partially derived from this dataset, as well as defined in the same surface space (fsLR) as the dataset.

### Compared with other approaches, the gwMRF model yielded parcellations with higher connectional homogeneity

Figure 3 compares the connectional homogeneity of parcellations generated with the gwMRF model with four publicly available parcellations. Using rs-fMRI data from the (A) GSP (N = 745) in fsaverage space, (B) HCP (N = 820) in fsLR space and (C) NKI (N = 205) in MNI space, connectional homogeneity was measured by averaging pairwise correlations within a parcel and across the cortex (details in *Methods*). We note that for the same parcellation, the number of parcels might be slightly different across coordinate systems due to masking and filtering operations (see *Methods*).

The gwMRF model generated parcellations that were significantly more connectionally homogeneous than Gordon (NKI: p = 9.4e-111, GSP: p ≈ 0), Craddock (NKI: p = 3.1e-59, GSP: p ≈ 0), Shen (NKI: p = 2.8e-28, GSP: p ≈ 0) and Glasser (NKI: p = 6.3e-110, GSP: p ≈ 0). No p value was computed for the HCP dataset because the HCP group-averaged dense connectivity matrix was utilized (so there was no access to connectional homogeneity computed in individual subjects). However, given that the HCP dataset was larger than the other two datasets, it is probably safe to assume that improvement in connectional homogeneity was statistically significant.

Across the datasets, the gwMRF model generated parcellations with average improvements of 8.0% (Gordon), 4.5% (Shen), 4.6% (Craddock) and 7.7% (Glasser). It is worth mentioning that the Glasser parcellation enjoyed inherent advantage for the HCP dataset because the Glasser parcellation was partially derived from this dataset and defined in the same surface space (fsLR) as the rs-fMRI data.

We also note that across all three datasets, the 333-area gwMRF parcellation was more connectionally homogeneous than the 360-area gwMRF parcellation even though more parcels should generally lead to higher homogeneity. This discrepancy arose because for the 333-area parcellation, border vertices were unlabeled to match the Gordon parcellation (see details in *Methods*), thus artificially boosting connectional homogeneity. Another interesting observation was that the Craddock parcellation enjoyed superior connectional homogeneity with respect to the Glasser parcellation in the GSP and NKI datasets even though the number of Craddock parcels was less than or equal to the number of Glasser parcels.

A final observation was that homogeneity computed in MNI space (Figure 3C) appeared significantly lower than homogeneity computed in surface spaces (Figures 3A and 3B), consistent with observations from previous studies (Zuo et al. 2013; Jiang and Zuo 2016).One major reason was probably because the NKI data was not smoothed, but another reason might be the significantly better registration accuracy achieved by a surface coordinate system (Fischl et al. 1999, 2008; Yeo et al. 2010). The surface coordinate system is a natural choice for this work because of our focus on the cerebral cortex. However, volumetric coordinate systems (e.g., MNI) are invaluable for analyzing subcortical structures.

### Stability of parcellations across GSP training and test sets

Cerebral cortex parcellations with 400, 600, 800 and 1000 parcels were estimated separately from the GSP training and test sets. Percentage overlaps between the training and test parcellations were 81%, 76%, 75% and 69% respectively, suggesting stability better than or equal to previous parcellations (e.g., 71% reported by Gordon et al. 2016). Visual inspection of parcellation boundaries suggests relatively good agreement between the training and testing parcellations (Figure S4), especially for parcellations with fewer parcels.

### Cerebral Cortex Parcellations of 1489 participants

Cerebral cortex parcellations with 400, 600, 800 and 1000 parcels were computed using the full GSP dataset (N = 1489) and shown in Figure S5. These parcellations were computed in fsaverage6 space, but the parcellations are publicly available in fsaverage, fsLR and MNI152 space (https://github.com/ThomasYeoLab/CBIG/tree/master/stable_projects/brain_parcellation/Scha_efer2018_LocalGlobal).

Figure 4 (first row) shows the 7-network and 17-network parcellations from Yeo et al. (2011). Figure 4 (second row) shows the 400-area cerebral cortex parcellation where the color of each parcel was assigned based on its spatial overlap with the original 7-network and 17-network parcellations. The 400-area parcellation were clustered (details in *Methods)* using a similar approach to Yeo et al. (2011), revealing 7 and 17 networks (Figure 4 last row) that were visually very similar to the original networks (Figures 4 first row).

Therefore the gwMRF cerebral cortex parcellations largely preserved the community or network structure of the original dataset, although there were some small differences. For example, the default network (red) in the 7-network solution (Figure 4 last row) comprised more precuneus parcels compared with the optimal assignment (Figure 4 second row). Similarly, the salience/ventral attention network (violet) in the 17-network solution (Figure 4 last row) comprised less precuneus parcels compared with the optimal assignment (Figure 4 second row). Similar results were obtained with the 600-area, 800-area and 1000-area parcellations (Figures S6, S7 and S8).

### Comparison of 400-area Parcellation with Architectonic and Visuotopic Areas

Figure 5 overlays parcels of the 400-area parcellation on the boundaries of histologically defined architectonic areas 3, 4, 2, hOc5 and 17 on the left cerebral cortical hemisphere (Fischl et al. 2008; Van Essen et al. 2012a). Other architectonic areas (including those on the right hemisphere) are shown in Figure S9.

There were generally good correspondences between parcellation boundaries and architectonic areas, especially for areas 3, 4, 2 and area 17 in both hemispheres. However, it was also clear that the parcellation fractionates primary areas into sub-regions. In the case of somatomotor areas 3, 4 and 2, the fractionation might correspond to somatotopic representations of different body parts, consistent with motor task activations (Figure 6) and previous functional connectivity parcellations (Yeo et al. 2011; Glasser et al. 2016). The parcellation also appeared to fractionate area 17 along the eccentricity axis, since the parcels were oriented orthogonal to the calcarine sulcus.

**Figure 6.**
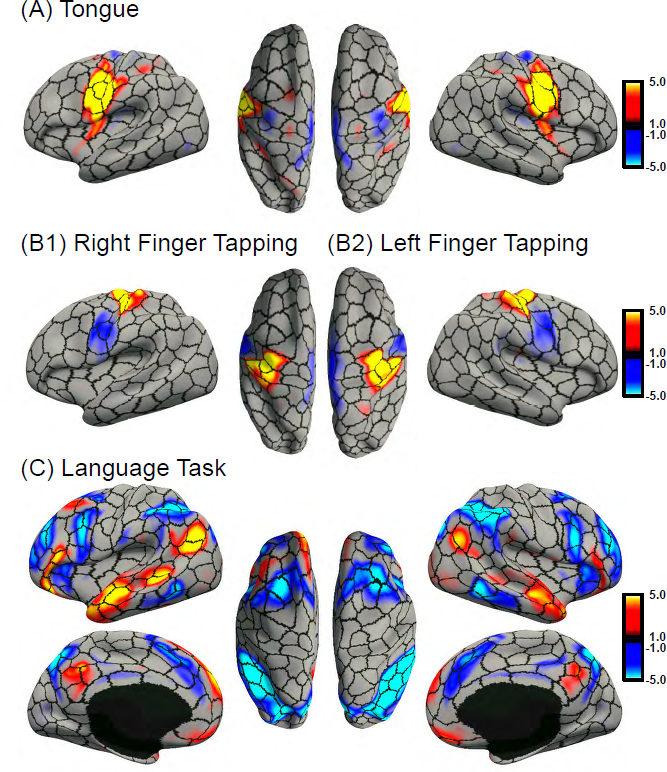
Group-average task activation maps of (A) tongue motion (tongue – average motor), (B1) right finger tapping (right finger – average motor), (B2) left finger tapping (left finger – average motor), and (D) language task (story – math) from the HCP dataset overlaid on boundaries (black) of 400-area cerebral cortex parcellation. Other task activation maps with different thresholds can be found in Figures S11A to S11E.

More muted correspondence with other architectonic areas was observed. For example, the parcel that maximally overlapped with area hOc5 appeared to extend beyond the architectonic boundaries in both hemispheres. Similarly, the parcels overlapping with area 18 appeared to extend beyond its dorsal boundary with area 19, but appeared to match well to its boundary with area 17 and its ventral boundary with area 19. The parcellation was unable to capture the dorsal aspect of area 1.

Figure S10 overlays boundaries of the 400-area parcellation on 18 visuotopically-mapped retinotopic areas (Abdollahi et al. 2014). There was generally good agreement between parcellation boundaries and retinotopic boundaries. This is especially true for retinotopic area V1 (which should theoretically correspond to architectonic area 17), which appeared to be further fractionated based on visual eccentricity. Other than the V1/V2 boundary, where there were corresponding parcel borders (ignoring the additional eccentricity boundaries), correspondences between parcel boundaries and other retinotopic regions were not immediately obvious. For example, multiple parcels spanned across both V2 and V3, which might reflect eccentricity organization within V2 and V3. Intriguingly, LO1 and LO2, which are in the same visual field map clusters (Wandell et al. 2007), appeared to be grouped together, although also fractionated, possibly also based on visual eccentricity organization (c.f. Figure 8C of Abdollahi et al., 2014). Overall, parcels in visual cortex might reflect supra-areal organization (Buckner and Yeo 2014) in addition to areal boundaries.

### Comparison of 400-area Parcellation with Task Activations and rs-fMRI

Figure 6 overlays the motor and language task activations from the HCP on the 400-area parcellation boundaries. The z-maps were averaged over 833 to 852 subjects (depending on the contrast) provided by the HCP (Barch et al. 2013). Other task contrasts with different thresholds can be found in Figures S11A to S11E. In general, a single contrast elicited task activations that cut across parcellation boundaries even with relatively high threshold (e.g., parietal portion of language contrast in Figure S11A). This is perhaps not surprising because a given task contrast likely involves a diverse set of cognitive processes supported by multiple cortical areas. Even if the task manipulation manages to elicit a single cognitive process, this cognitive process could be implemented across numerous cortical areas (Poldrack 2006; Barrett and Satpute 2013; Yeo et al. 2015a).

Figure S12 shows the functional inhomogeneity of individual parcels of the 400-area parcellation for different task contrasts. Visual inspection of Figures S11 and S12 suggests a complex relationship between the magnitude of activation (Figure S11) and functional inhomogeneity (Figure S12). To quantify this phenomenon, scatterplots of parcel activation magnitude (absolute value of average z-scores) against parcel functional inhomogeneity (Figure S13) suggest that higher activation magnitude was correlated with functional inhomogeneity. Therefore functional inhomogeneity (Figures 2, S12) is mostly driven by strong activation or de-activation.

Figure 7 shows the connectional homogeneity of each parcel of the 400-area parcellation in the GSP and HCP datasets. Although the gwMRF model achieved better connectional homogeneity than other parcellations (Figure 3), not all parcels were equally homogeneous (Figure 7). The spatial variation in homogeneity was consistent across the GSP and HCP datasets despite differences in scanners, acquisition protocols and preprocessing strategies. More specifically, parcels in the ventral temporal lobe and orbital frontal cortex were of lower homogeneity, probably due to lower signal-to-noise ratio (SNR) in those regions.

**Figure 7.**
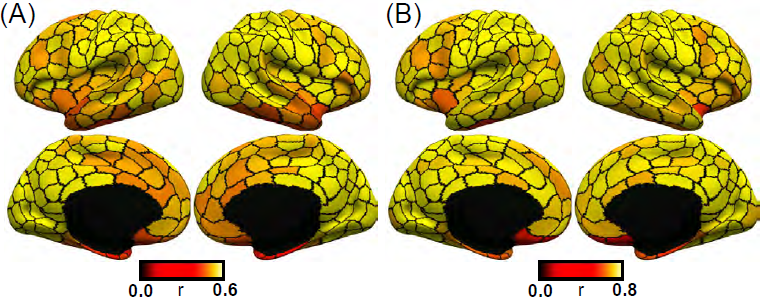
Functional connectivity homogeneity of each parcel of 400-area cerebral cortex parcellation based on (A) GSP and (B) HCP datasets. Overall homogeneity for the GSP and HCP datasets were 0.58 and 0.78 respectively. Note that the parcellation was derived from the GSP dataset. However, acquisition and processing differences might lead to higher homogeneity in the HCP dataset.

Connectional homogeneity was higher in the HCP dataset (0.78) compared with the GSP dataset (0.58), even though the parcellation was derived from the GSP dataset. The difference might arise from the multiband protocol and longer acquisition time in the HCP data, which might result in higher SNR and therefore connectional homogeneity. In addition, the HCP utilized a function-based inter-subject registration algorithm (Robinson et al. 2014), which might reduce inter-subject variability and therefore increased connectional homogeneity. Finally, the HCP preprocessing utilized a Wishart roll-off denoising, which might implicitly introduce more smoothing than the 6mm fwhm smoothing applied to the GSP data.

### Volume distribution of the 400-area parcellation

Figure S14 illustrates distributions of parcel volume of the 400-area parcellation, and four previously published parcellations (Craddock, Shen, Glasser, Gordon). The volumetric ratios of the largest parcel to the smallest parcel were 16 (gwMRF), 3.8 (Craddock), 3.8 (Shen), 45 (Glasser) and 443 (Gordon). The volumetric ratios of the 90th percentile parcel to the 10th percentile parcel were 3.2 (gwMRF), 1.7 (Craddock), 1.6 (Shen), 5.7 (Glasser) and 8.3 (Gordon).

## Discussion

We developed a gradient-weighted Markov Random Field (gwMRF) model that integrated the local gradient and global similarity approaches to brain parcellation. Cerebral cortical parcellations generated by the gwMRF model were compared with three rs-fMRI parcellations derived from fully automatic algorithms (Craddock et al., 2012, Shen et al., 2013; Gordon et al., 2016) and a multimodal parcellation derived from a semi-automatic method (Glasser et al., 2016). Compared with the fully automatic parcellations, the gwMRF parcellations had similar agreement with the boundaries of architectonic and visuotopic areas. Compared with the Glasser parcellation, the gwMRF parcellation had worse alignment with architectonic and visuotopic boundaries, which might reflect manual selection of multimodal information by an anatomist in the Glasser parcellation to match prior knowledge of areal boundaries. Parcellations generated by the gwMRF model enjoyed superior functional and connectional homogeneity compared to all other parcellations. Furthermore, the parcellations recapitulated the network structure of the original vertex-wise fMRI data.

### Fusion model generates parcellations that align with architectonic areas

The local gradient approach for delineating cortical areas with rs-fMRI was first proposed by the pioneering work of Cohen et al. (2008). Although the approach has been largely applied to rs-fMRI (Nelson et al. 2010; Hirose et al. 2012; Wig et al. 2014b; Laumann et al. 2015; Gordon et al. 2016; Xu et al. 2016), recent work has also applied this approach to multimodal data (Glasser et al., 2016). It has been suggested that the gradient approach is uniquely suited for delineating histologically defined cortical areas (Wig et al., 2014b; Gordon et al., 2016) compared with global clustering approaches (e.g., mixture models, spectral clustering).

More specifically, an influential paper (Wig et al., 2014b) demonstrated that the local gradient approach was able to delineate area 17 unlike global approaches, such as the infomap algorithm (Power et al., 2011) or the mixture model (Yeo et al., 2011). However, the comparison was imperfect because both Power et al. (2011) and Yeo et al. (2011) parcellated the cerebral cortex into less than twenty spatially distributed networks, corresponding to roughly one hundred parcels. Since there might be 300 to 400 human cortical areas based on extrapolation from monkey data (Van Essen et al. 2012a), it should not be surprising that relatively low resolution parcellations (Power et al., 2011; Yeo et al., 2011) were unable to fully differentiate cortical areas.

Indeed our experiments suggested that the global similarity approach could produce parcellation boundaries that closely match the boundaries of area 17. On the other hand, the global similarity approach was unable to differentiate between areas 3 and 4 because of strong RSFC between the two areas (Figure 1). This strong functional connectivity might result from signal bleeding across the central sulcus, but might also have its origins in strong anatomical connections between the primary somatosensory and motor areas (Jones 1986). On the other hand, the local gradient approach was unable to capture the boundaries of area 44 unlike the global similarity approach (Figure 1).

Based on visual inspection, gwMRF parcellations obtained by fusing both local gradient and global similarity approaches were able to capture architectonic boundaries better than either approach (Figures 1, 5 & S9), although the numerical results are not statistically different (Figure S2). Overall, parcellations generated by the gwMRF fusion model had comparable architectonic alignment with parcellations derived from fully automatic approaches (Craddock et al. 2012; Shen et al. 2013; Gordon et al. 2016). The semi-automatic parcellation (Glasser et al., 2016) had better alignment with boundaries of areas 17, 18, hOc5, 3, 4, but worse alignment with boundaries of areas 2, 6, 44 and 45. The variable differences might reflect priorities of the anatomist who manually selected among competing multimodal information.

### Fusion model generates parcellations that align well with visuotopic areas

Alignment with visuotopically defined retinotopic areas (Abdollahi et al., 2014) was considered. As illustrated in Figure 5, parcellations generated by the gwMRF model have comparable visuotopic alignment with parcellations from fully automatic approaches (Craddock et al. 2012; Shen et al. 2013; Gordon et al. 2016), but worse alignment than the semi-automatic multimodal approach (Glasser et al., 2016).

Visual inspection (Figure S8) suggested that the parcellation boundaries were reasonably aligned with retinotopic boundaries, especially V1. However, there is generally no one-to-one mapping between parcels and retinotopic areas. For example, V1 appeared to be further fractionated based on visual eccentricity. This fractionation likely extended to other retinotopic areas in the visual cortex, suggesting that parcels in visual cortex might reflect supra-areal (e.g., visual eccentricity) organization in addition to areal boundaries (Buckner and Yeo, 2014).

It is worth noting that the retinotopic and architectonic boundaries were themselves not in complete alignment. For example, architectonic area 18 did not seem to correspond well to retinotopic area V2 or V3. Similarly hOc5 appeared to correspond to multiple retinotopic areas (MT and pMST). Therefore we did not expect perfect matching between rs-fMRI parcellation boundaries with architectonic and retinotopic boundaries. Nevertheless, the performance of the multimodal parcellation (Glasser et al., 2016) suggested significant room for improvements.

### Fusion model generates parcellations that are functionally homogeneous

Functional homogeneity of a parcellation was evaluated using task-fMRI. One important consideration was the choice of task contrast to evaluate functional homogeneity. A stricter contrast might provide access to a purer cognitive construct, but it is unlikely any of the HCP task contrast tapped into a single cognitive function that activated only a single cortical area. Previous work has approached this issue by evaluating the fraction of task-activated vertices that fell within rs-fMRI parcels as opposed to the boundaries between parcels (Laumann et al. 2015). Their approach made sense in the context of their implementation of the local gradient approach, which introduced non-uniformly thick borders between rs-fMRI parcels (Gordon et al., 2016). Because there was a need to threshold the task activation, multiple thresholds were considered (Laumann et al. 2015).

Here we avoided the need to threshold task activation by simply computing the standard deviation of task activation z-values within each parcel (Figure 2). A lower standard deviation would indicate a parcel is more functionally homogeneous. This approach did not require the task contrast to only recruit a single cognitive function or activate a single cortical area. Instead, high functional homogeneity could be achieved as long as activation strength was uniform within each cortical area, which was reasonable since we were trying to isolate neurobiological atoms within the cerebral cortex. If task activations were not uniform within a parcel, this would imply functional heterogeneity within the parcel, suggesting that the parcel was not a neurobiological atom. Since the functional inhomogeneity measure did not require the task contrast to isolate a single cognitive process, we decided to utilize all contrasts across all seven cognitive domains, rather than having to justify certain contrasts over others (Figure 2).

Given the strong link between task-fMRI and rs-fMRI (Smith et al. 2009; Mennes et al. 2010; Cole et al. 2014; Krienen et al. 2014; Bertolero et al. 2015; Yeo et al. 2015a; Tavor et al. 2016), and since the gwMRF model produced parcellations that were the most connectionally homogeneous (previous section), it was perhaps not surprising that the gwMRF parcellations were also the most functionally homogeneous (Figure 2). Like in the case of connectional homogeneity, we note that the HCP task-fMRI data were collected from a different scanner with a different acquisition protocol and processed in a different way from the GSP data used to derive the gwMRF parcellations.

Intriguingly, functional inhomogeneity varied across cognitive domains (Figure 2) with the working memory task exhibiting greater inhomogeneity than the other tasks. Possible mundane reasons might include variation in task design and overall task duration across cognitive domains. However, the relatively larger inhomogeneity of the working memory task was consistent with previous work exploring the areal organization of a highly sampled individual (see Figure 3 of Laumann et al., 2015).

### Fusion model generates parcellations that are connectionally homogeneous

One potential use of rs-fMRI parcellations is for reducing the dimensionality of fMRI data in future studies (Eickhoff et al., 2017). For example, it is common to average the fMRI time courses within each parcel of a brain parcellation and use the resulting average time courses to compute functional connectivity matrices for studying mental disorders (Fair et al. 2013; Baker et al. 2014), individual subject differences (Finn et al. 2015; Yeo et al. 2015b), graph theoretic analyses (Salvador et al. 2005; Supekar et al. 2008; Bullmore and Sporns 2009; Meunier et al. 2009; Guye et al. 2010; He and Evans 2010; Wang et al. 2010; Zalesky et al. 2010a; Fornito et al. 2013) or neural mass modeling (Ghosh et al. 2008; Honey et al. 2009; Deco et al. 2013; Zalesky et al. 2014; Betzel et al. 2016). For the dimensionality reduction to be meaningful, the representative time course of a parcel should be similar to all time courses within the parcel (Zuo and Xing 2014; Eickhoff et al., 2017). Our results suggest that gwMRF parcellations exhibit significantly better functional connectivity homogeneity than parcellations derived from local gradient (Gordon et al., 2016; Glasser et al., 2016) or global similarity (Craddock et al., 2012; Shen et al., 2013) approaches (Figure 3). Therefore the gwMRF parcellations might be a useful dimensionality reduction tool for the fMRI community.

We note that the gwMRF parcellations were derived from the GSP data acquired from Siemens Tim Trio scanners and preprocessed using a pipeline involving whole brain signal regression in FreeSurfer fsaverage space (Holmes et al. 2015). The improvement in functional connectivity homogeneity (Figure 3) was replicated in datasets collected from different scanners and acquisition protocols (HCP and NKI), preprocessing pipelines (ICA-FIX and CompCor) and coordinate systems (fsLR and MNI). Furthermore, while the GSP and HCP datasets consisted of young healthy adults, the NKI dataset consisted of participants ranging from ages 6 to 85, suggesting that the gwMRF parcellations generalized well to different populations, and were suited for studies across different stages of the human lifespan.

Finally, even though connectional homogeneity was high, there was spatial variation in homogeneity (Figure 7) with parcels in low SNR regions (ventral temporal lobe and orbitofrontal cortex) being of lower homogeneity. The lower SNR might potentially decrease parcellation accuracy within these regions. Future work might explore if post-hoc correction of reliability and SNR could improve parcellation quality (Mueller et al. 2015).

### Towards multi-feature and multimodal parcellations

Despite the generally good match between the gwMRF parcellations and boundaries of architectonic and retinotopic areas, there was significant room for improvement. For example, the 400-area parcellation did not conform well to the boundaries of architectonic area hOc5, as well as the boundaries between areas 3a and 3b or areas 4a and 4p. Similarly, the 400-area parcellation did not conform well to many retinotopic boundaries but might instead reflect supra-areal (e.g., visual eccentricity) organization. These results suggested that the fusion of local gradient and global similarity approaches to rs-fMRI was insufficiently sensitive to the boundaries of certain cortical areas.

However, rs-fMRI might not necessarily be intrinsically insensitive to these boundaries. For example, a recently developed rs-fMRI approach (Glasser et al. 2016) exploited the fact that cortical regions representing the same visual field locations are anatomically (Cragg 1969; Zeki 1969; Van Essen and Zeki 1978; Maunsell and Van Essen 1983) and functionally (Yeo et al. 2011; Wang et al. 2012; Striem-Amit et al. 2015) connected. This elegant approach (together with manual intervention) allowed the exquisite delineation of visual areas, resulting in better agreement with retinotopic boundaries than gwMRF parcellations (Figure S3). Future work could potentially integrate this topographic rs-fMRI method with approaches used in the current work.

Contributions from other modalities might be necessary (Toga et al., 2006; Eickhoff et al., 2011; Bzdok et al., 2013; Wang et al., 2015a; Glasser et al., 2016) if there were certain biological boundaries where rs-fMRI is truly intrinsically insensitive. For example, the relative myelin mapping technique allowed beautiful delineation of somatomotor areas 4a, 4p, 3a, 3b, 1 and 2 (Glasser and Van Essen 2011; Glasser et al. 2016), while the boundaries between areas 4a and 4p, as well as between areas 3a and 3b, were not visible with the standard application of local gradient to RSFC (see Figure 2 of Gordon et al., 2016).

Therefore fusion of myelin mapping with rs-fMRI might potentially improve brain parcellation.

One challenge of multimodal integration is the resolution of multimodal conflicts. In the seminal semi-automatic multimodal parcellation (Glasser et al., 2016), the anatomist explicitly ignored strong RSFC gradients within somatomotor cortex and strong intra-V1 gradients based on prior neuroanatomical knowledge about sensory and motor cortical areas. However, while prior knowledge is abundant and reliable for sensory-motor cortex, prior knowledge is weak for the human association cortex, which has undergone significant evolutionary expansion and changes over the millions of years of evolution separating humans from macaques (Preuss 2004; Van Essen and Dierker 2007; Hill et al. 2010; Buckner and Krienen 2013). As such it is unclear how well the accuracy of Glasser parcellation (which was extremely impressive in sensory-motor cortex as seen in Figures S2D and S3) extended to association cortex (Yeo and Eickhoff, 2016). Indeed, the advantage of the Glasser parcellation largely disappeared for cortical areas 44 and 45 (Figure S2D), which are thought to be involved in language processing (Nishitani et al. 2005; Davis et al. 2008). Consequently, there is a need to develop methods that can automatically select among competing gradients. An advantage of utilizing fully automatic methods is that good parcellation performance within sensory-motor cortex might be assumed to extend to association cortex, assuming no fundamental difference between sensory-motor and association cortices, which might not necessarily be true (Buckner and Krienen 2013).

There are other costs to trading off between conflicting modalities. For example, while the Glasser parcellation achieved extremely good alignment with architectonic and visuotopic boundaries (Figures 2 and 5), it had significantly worse functional and connectional homogeneity (Figures 3 and 4), even though the Glasser parcellation was partially derived from the HCP task-fMRI and rs-fMRI used to evaluate functional and connectional homogeneity. On average, the Glasser parcellation exhibited 7.7% worse connectional homogeneity than the gwMRF parcellation and even had worse connectional homogeneity than the Craddock parcellation in the GSP and NKI datasets (Figure 4).

The reason was that our comparisons controlled for the number of parcels and therefore the Glasser parcellation “wasted” precious parcels distinguishing cortical areas with indistinguishable RSFC. For example, the Glasser parcellation differentiated between areas 3a and 3b, even though the tongue representations in areas 3a and 3b had very similar rs-fMRI and task-fMRI properties. On the other hand, by treating area 3a as a single parcel and area 3b as a single parcel, the Glasser parcellation “lost” the opportunity to differentiate among body representations (e.g., hand and tongue) with distinctive rs-fMRI and task-fMRI properties.

We note that there are functional differences between areas 3a and 3b – area 3b is thought to be involved in somatic sensation (Iwamura 1998), while area 3a is thought to be involved in proprioception (Krubitzer et al. 2004). However, given that there are only subtle differences between the two areas as measured by rs-fMRI and task-fMRI, differentiating the two areas does not seem very useful if the goal is to reduce the dimensionality of fMRI data. Conversely, differentiating the hand and tongue representations would be extremely useful when modeling behavioral tasks with button presses. Further discussion of this issue can be found in Eickhoff et al. (2017).

### Shape, size and lateralization of parcels

Compared with local gradient approaches (Gordon et al., 2016; Glasser et al., 2016), global similarity approaches (Craddock et al., 2012; Shen et al., 2013) appeared to generate rounder parcels (Figure S1) with more uniform sizes (Figure S13). One possible reason is that global similarity approaches are explicitly optimizing for homogeneous parcels and are therefore more sensitive to fMRI smoothness. Another possible reason is that global similarity approaches (Craddock et al., 2012; Shen et al., 2013; Ryali et al., 2013; Honnorat et al., 2015) often require an additional regularization objective to ensure spatially connected parcels, although it is worth noting that local gradient approaches implicitly constrain parcels to be spatially connected, such as by the use of the watershed algorithm (Gordon et al., 2016). By fusing local gradient and global similarity approaches, and by weakening the spatial connectedness objective during the optimization procedure, the gwMRF model generated parcels of intermediate roundness and size distribution (Figures S1, S13).

From the neuroscience perspective, traditionally defined cortical areas can be quite irregular (Van Essen et al., 2012a). In terms of size, Van Essen et al. (2012a) found that the ratio of the largest and smallest cortical areas was roughly 200, which is closest to the Gordon parcellation. In terms of shape, cortical areas can range from relatively round areas (e.g., area 44) to narrow, long areas (e.g., area 3). The 400-area gwMRF parcellation split V1 into visual eccentricity bands and areas 3 into somatotopic sub-regions. While these splits were biologically meaningful, they also increased parcel uniformity. Since V1 is one of the largest cortical areas (Van Essen et al., 2012a), splitting V1 resulted in parcels with more uniform sizes. Similarly, splitting the long and thin cortical area 3 into somatotopic sub-regions resulted in parcels that were rounder. However, this explanation might not extend to non-topographically organized cortical areas.

Given that gwMRF and Gordon parcellations achieved similar alignment with architectonic and visuotopic boundaries, gwMRF parcellations might represent the spatial layout of traditional cortical areal boundaries almost as well as gradient based approaches. On the other hand, given that gwMRF parcellations enjoyed superior functional and connectional homogeneity, gwMRF parcellations might be more useful than gradient-based parcellations when utilized as a dimensionality reduction tool for new fMRI data.

Finally, the gwMRF parcellations were derived for each hemisphere separately, although the local RSFC gradients were computed based on whole brain RSFC. Because of the spatial connectedness term in the gwMRF objective function, applying the procedure to both hemispheres should in theory yield similar parcellations. While functional systems (e.g., default and dorsal attentional networks) are known to span across both hemispheres, well-known functional lateralization within the cerebral cortex (e.g., Corbetta and Shulman, 2002) also suggests that a bilateral functional system might include an asymmetric distribution of parcels (Figure 4). Indeed, visual inspection of the local RSFC gradients (Figure 2 of Gordon et al., 2016) suggested that certain gradients were present in one hemisphere but not the other. Consequently, the gwMRF and Gordon parcellations were both asymmetric, consistent with most parcellation procedures in the literature (Craddock et al., 2012; Shen et al., 2013; Honnorat et al., 2015). One major exception was the recent multimodal parcellation, where the anatomists explicitly ignored asymmetric gradient information (Glasser et al., 2016).

### “Optimal” resolution of brain parcellation

A number of metrics has been proposed in the connectivity based parcellation literature to estimate or justify the number of brain parcels (Yeo et al. 2011; Eickhoff et al. 2015). There is also a wide range of machine learning techniques that seek to estimate the number of modules in a clustering problem, including Bayesian information criterion (BIC; Fraley & Raftery 1998) and stability analysis (Lange et al. 2004). However, these approaches are unlikely to estimate a truly optimal number of clusters because of approximations necessary to compute the metrics (Fraley and Raftery 1998). In many cases (e.g., stability analysis), the estimated number of clusters might partially reflect the size of the dataset, reducing confidence that the estimation reflected biology.

Furthermore the brain is hierarchically organized (Churchland and Sejnowski 1988) from molecules (1Å) to synapses (1μm) to neurons (100μm) to areas (1cm) to systems (10cm). Even at the millimeter resolution of MRI, hierarchical organization can be observed. For example, large-scale networks (e.g., default, dorsal attention, etc) comprise multiple cortical areas. Given the heterogeneity of cortical areas (Kaas 1987; Amunts and Zilles, 2015), cortical areas can be further subdivided. For example, the primary motor area can be divided into distinct body representations, such as the hand and mouth. The hand representation can in turn be subdivided into wrist and individual fingers.

Consequently, it is unclear if there is a correct number of brain parcels or if cortical areas are necessarily the optimal resolution for cortical parcellations. For example, Glasser et al. (2016) and Gordon et al. (2016) extracted V1 as a single parcel, while our 400-area parcellation divided V1 into parcels based on their visual eccentricity. Similarly, Gordon et al. (2016) and our 400-area parcellation divided somatomotor cortex into parcels, presumably based on body representations. The different portions of area 3a represent distinct body parts and might therefore be considered as distinct computational units (Yeo and Eickhoff, 2016). As such, one could argue that separating the different somatotopic regions might be useful for future computational modeling. For example, it might be useful to differentiate between the hand and foot motor regions when modeling a behavioral task with button presses. As an additional example, differentiating between central and peripheral V1 might be useful when modeling a task involving peripheral visual distractors.

Furthermore, one might expect that different resolution parcellations might be useful for different applications. For example, if the effect of interest is highly focal (or diffuse), then a higher (or lower) resolution parcellation might be more effective (Jones et al. 2005; Zalesky et al. 2010b). Therefore we generated cortical parcellations consisting of 400, 600, 800 or 1000 parcels.

### Confounds when comparing brain parcellations

Comparing brain parcellations generated from different labs is difficult. A major confounding factor is the number of parcels. One approach to mitigating this issue is the use of a null model to statistically test whether a brain parcellation performs better than random parcellations with the same number of parcels, as well as distribution of parcel sizes and shapes (Gordon et al. 2016). Here we employed the more explicit approach of matching the number of parcels when comparing parcellations.

Another major confounding factor not widely acknowledged in the literature is the comparisons of parcellations generated from different data or coordinate systems. Here we employed multimodal data from multiple scanners, acquisition protocols and preprocessing strategies across different coordinate systems, thus increasing confidence that our parcellations would generalize well to data collected from a different scanner preprocessed in a different way.

Projecting a parcellation to a coordinate system different from where it was derived can significantly deteriorate the parcellation quality. For example, parcellations generated by our fusion model achieved dramatically better connectional homogeneity compared with the Shen (7.6%) and Craddock (7.2%) parcellations in fsaverage space. The improvements persisted but decreased to 1.0% and 1.8% respectively in MNI space.

Finally, our comparisons were largely limited to three publicly available rs-fMRI parcellations (Craddock et al., 2012; Shen et al., 2013; Gordon et al., 2016) and one multimodal parcellation (Glasser et al., 2016). More comprehensive comparisons with other approaches (Arslan et al., 2017), as well as parcellations from different modalities, such as diffusion MRI (Fan et al., 2016) or histology (Eickhoff et al., 2005; Ding et al., 2016) are left to future work.

### Limitations and future work

The cerebral cortex forms spatially organized connections with subcortical structures (Jones 1985; Haber 2003; Strick et al. 2009). Here, we limited our parcellations to the cerebral cortex although the gradient maps we utilized took into account cortico-subcortical connectivity (Gordon et al., 2016). While our approach is in principle applicable to subcortical structures, accurate parcellation of the entire brain in a single step is non-trivial because of significant SNR difference between the cerebral cortex and subcortical structures. As such, subcortical structures might be more effectively parcellated separate from the cerebral cortex, as is performed in many studies (Di Martino et al., 2008; Zhang et al., 2008; Krienen and Buckner, 2009; Buckner et al., 2011; Choi et al., 2012; Dobromyslin et al., 2012; but see Craddock et al., 2012; Shen et al., 2013). For example, fMRI signals from the adjacent visual cortex tend to bleed into adjacent cerebellar regions that form close anatomical loops with the motor cortex, but not the visual cortex (Strick 1985; Schmahmann and Pandya 1997). Therefore additional regression steps were necessary to remove these confounds (Buckner et al., 2011).

The cerebral cortical parcellations we derived were obtained by averaging data across many participants. Given well-known individual differences in brain functional organization (Mueller et al. 2013; Laumann et al. 2015; Glasser et al. 2016; Gordon et al. 2017), the group level parcellations derived in this work might not be an optimal fit to individual subjects. The issue might be exacerbated for higher-resolution parcellations (e.g., 1000-area parcellation) because the smaller group-level parcels are more likely to be mismatched to the intrinsic organization of individual subjects. Therefore subject-specific brain network (Hacker et al. 2013; Gordon et al. 2016; Harrison et al. 2015; Varoquaux et al., 2011; Wang et al. 2015b) or areal estimation (Laumann et al. 2015; Glasser et al. 2016) might be potentially important for understanding individual differences in behavior and disorder. The extension of the gwMRF model to individual subjects is left for future work.

### Conclusion

We developed an rs-fMRI parcellation algorithm that exploited local gradients in resting-state functional connectivity, while maximizing similarity of rs-fMRI time courses within a parcel. The resulting cerebral cortex parcellations were functionally and connectionally homogeneous, and were in good agreement with certain architectonic and visuotopic boundaries. Multi-resolution parcellations are provided as reference atlases for future characterization with other modalities, graph theoretic analysis and neural mass modeling (https://github.com/ThomasYeoLab/CBIG/tree/master/stable_projects/brain_parcellation/Scha_efer2018_LocalGlobal).

## Acknowledgements

This work was supported by Singapore MOE Tier 2 (MOE2014-T2-2-016), NUS Strategic Research (DPRT/944/09/14), NUS SOM Aspiration Fund (R185000271720), Singapore NMRC (CBRG/0088/2015), NUS YIA and the Singapore National Research Foundation (NRF) Fellowship (Class of 2017). Alexander Schaefer was supported by a DAAD postdoctoral fellowship. Timothy Laumann was supported by NIMH (MH100872). Xi-Nian Zuo acknowledged support from the National Basic Research (973) Program (2015CB351702) and the Natural Science Foundation of China (81471740, 81220108014). Avram J Holmes acknowledged support from NIMH (K01MH099232). The research also utilized resources provided by the Center for Functional Neuroimaging Technologies, P41EB015896 and instruments supported by S10RR023401 and S10RR023043 from the Athinoula A. Martinos Center for Biomedical Imaging at the Massachusetts General Hospital. Data were provided [in part] by the Human Connectome Project, WU-Minn Consortium (Principal Investigators: David Van Essen and Kamil Ugurbil; 1U54MH091657) funded by the 16 NIH Institutes and Centers that support the NIH Blueprint for Neuroscience Research; and by the McDonnell Center for Systems Neuroscience at Washington University.

## Supplemental Figures

**Figure S1A.**
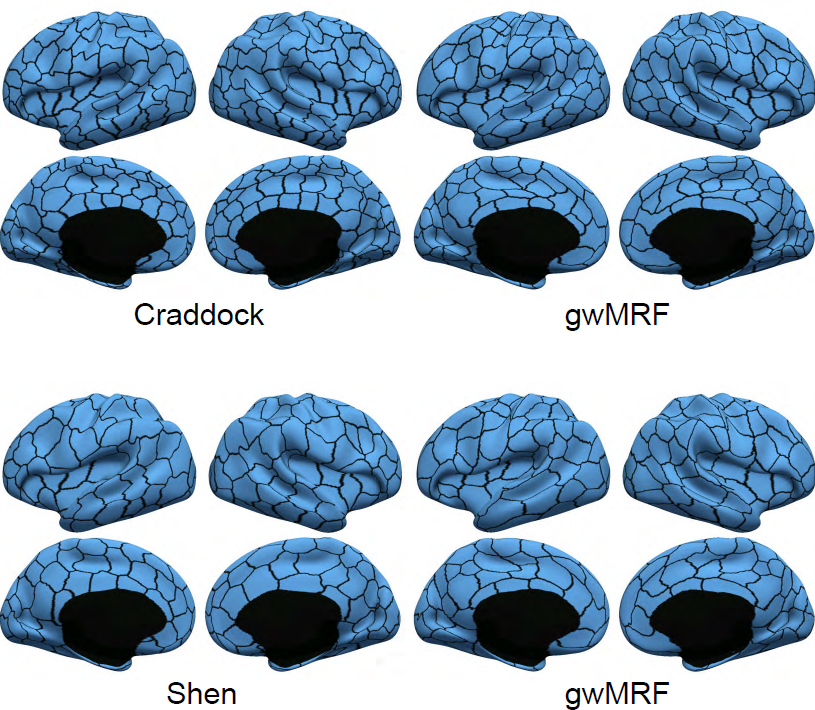
Craddock and Shen parcellations (left) and gwMRF parcellations with matching number of parcels (right).

**Figure S1B.**
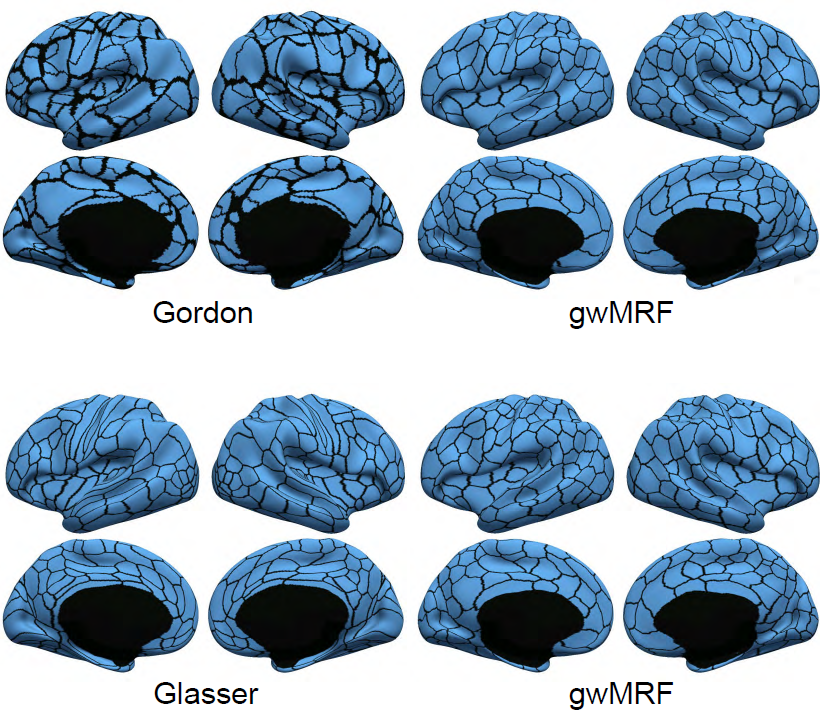
Gordon and Glasser parcellations (left) and gwMRF parcellations with matching number of parcels (right).

**Figure S2.**
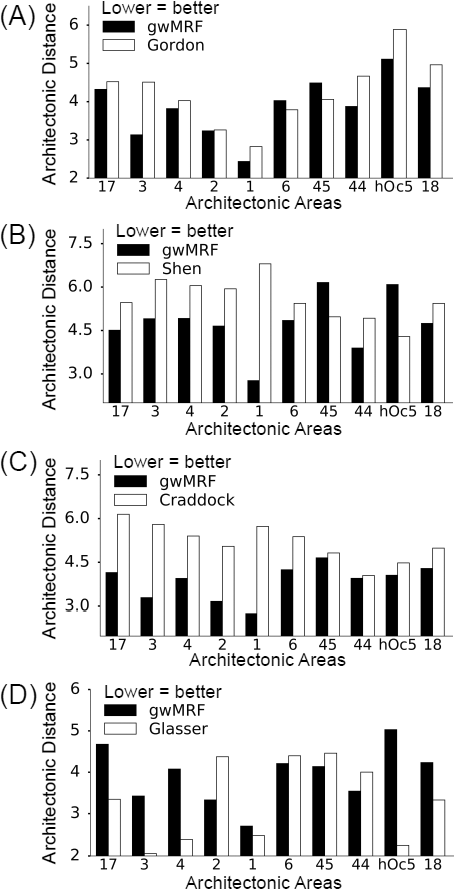
Distance to architectonic boundaries as measured by average Hausdorff distance (mm). Lower distance indicates better agreement. The gwMRF fusion approach generated parcellations that achieved (A) comparable architectonic distance with Gordon (p = 0.076 uncorrected), (B) better distance than Shen (p = 2.7e-5 uncorrected), (C) better distance than Craddock (p = 0.049 uncorrected), and (D) worse distance than Glasser (p = 0.042 uncorrected). It is worth mentioning that the Glasser parcellation required an anatomist to manually select specific multi-modal information matching prior knowledge of cortical areas. We note that the parcellations comprised 333, 348, 236 and 360 parcels in subplots A, B, C and D respectively. Therefore comparisons between subplots are not meaningful because more parcels lead to more boundary vertices, and therefore lower geodesic distances (on average).

**Figure S3.**
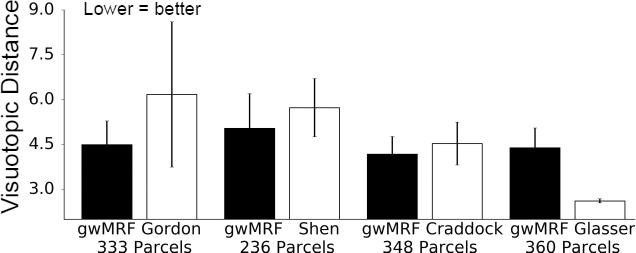
Distance to visuotopic boundaries (Abdollahi et al., 2014) as measured by average Hausdorff distance (mm). Lower distance indicates better agreement. The gwMRF parcellations achieve (A) lower visuotopic distance than Gordon (p = 8.7e-3), (B) similar distance to Craddock (p = 0.37), (C) similar distance to Shen (p = 0.21), and (D) worse distance than Glasser (p = 1.8e-6). It is worth mentioning that the Glasser parcellation required an anatomist to manually select specific multi¬modal information matching prior knowledge of cortical areas. Like before, comparisons between the subplots are not meaningful because the number of parcels is different across the parcellations.

**Figure S4.**
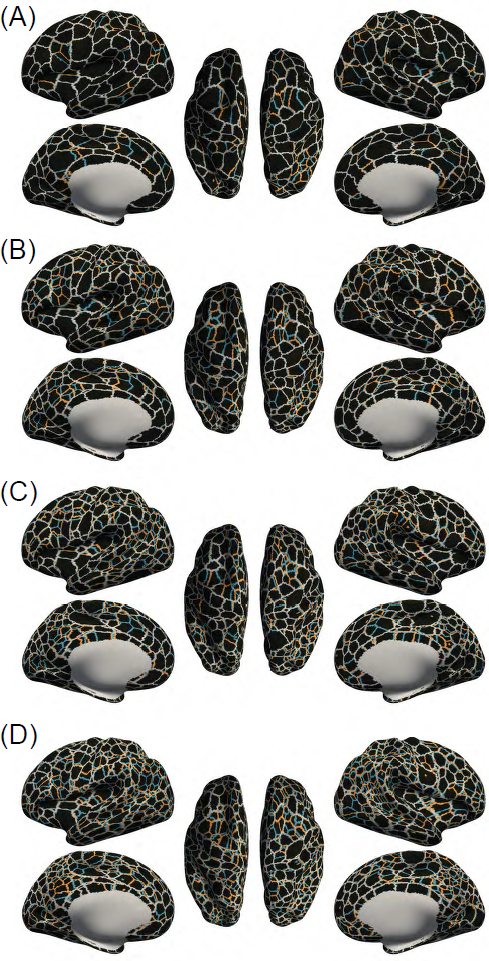
Agreement between gwMRF parcellations estimated from GSP training and test sets. (A) 400-area, (B) 600-area, (C) 800-area, (D) 1000-area. Red lines indicate boundaries of GSP training set parcellation. Blue lines indicate boundaries of GSP test set parcellation. White lines indicate overlapping parcellation boundaries between the two datasets.

**Figure S5.**
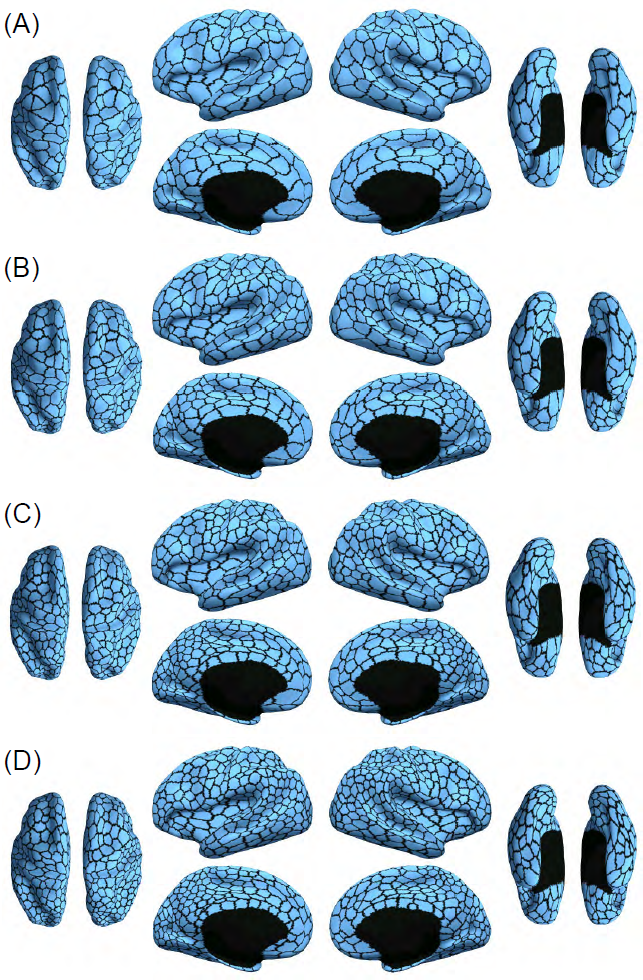
Cerebral cortex parcellations with (A) 400 (B) 600 (C) 800 and (D) 1000 parcels based on the entire GSP dataset of 1489 subjects.

**Figure S6.**
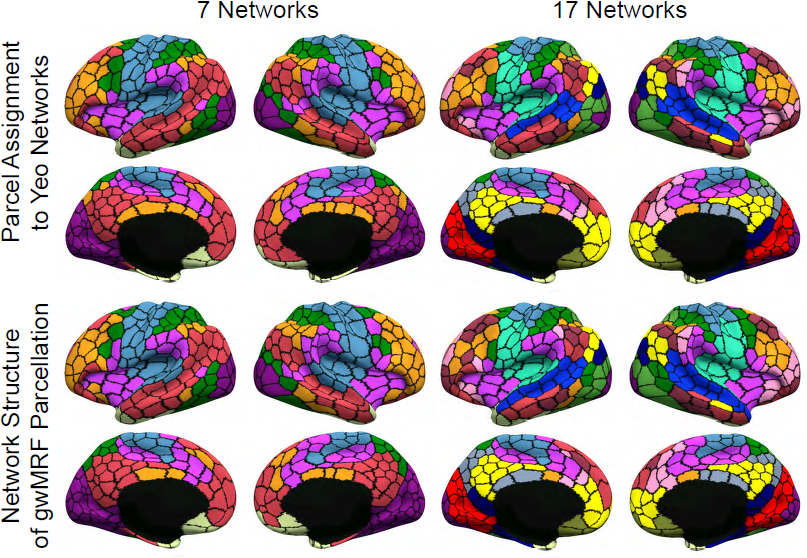
Network structure is preserved in the 600-area parcellation. First row shows each gwMRF parcel assigned a network color based on spatial overlap with networks from Yeo et al. (2011). Second row shows community structure of gwMRF parcellation after clustering. Observe striking similarity between first and second rows, suggesting that network structure of the original resolution data is preserved in the 600-area parcellation.

**Figure S7.**
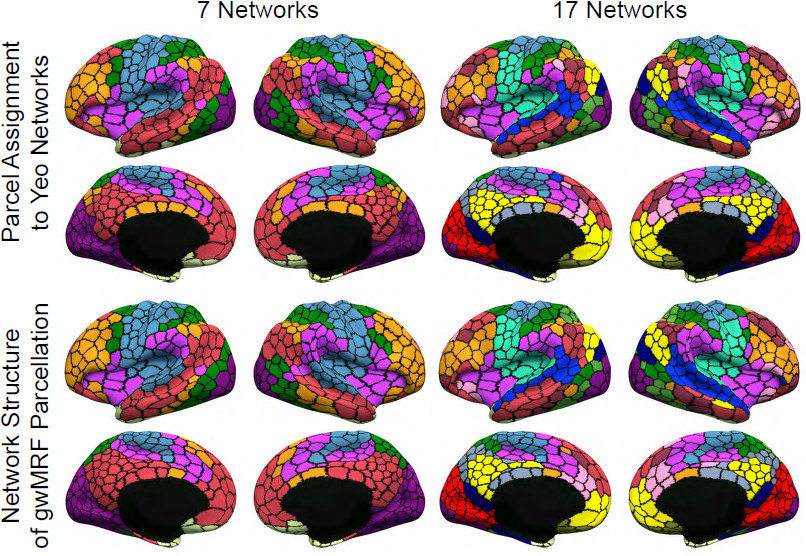
Network structure is preserved in the 800-area parcellation. First row shows each gwMRF parcel assigned a network color based on spatial overlap with networks from Yeo et al. (2011). Second row shows community structure of gwMRF parcellation after clustering. Observe striking similarity between first and second rows, suggesting that network structure of the original resolution data is preserved in the 800-area parcellation.

**Figure S8.**
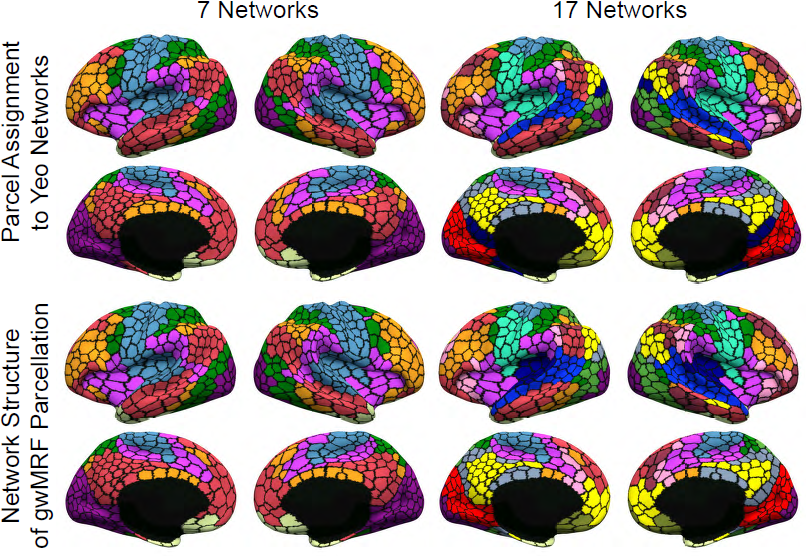
Network structure is preserved in the 1000-area parcellation. First row shows each gwMRF parcel assigned a network color based on spatial overlap with networks from Yeo et al. (2011). Second row shows community structure of gwMRF parcellation after clustering. Observe striking similarity between first and second rows, suggesting that network structure of the original resolution data is preserved in the 1000-area parcellation.

**Figure S9A.**
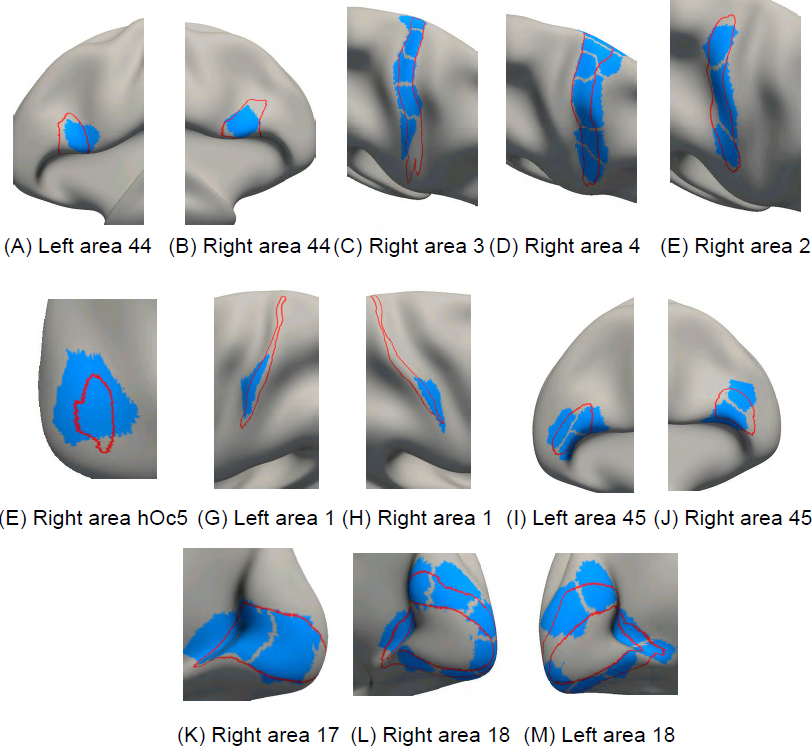
Parcels (blue) of the 400-area cerebral cortex parcellation overlaid on (red) boundaries of histologically defined areas

**Figure S9B.**
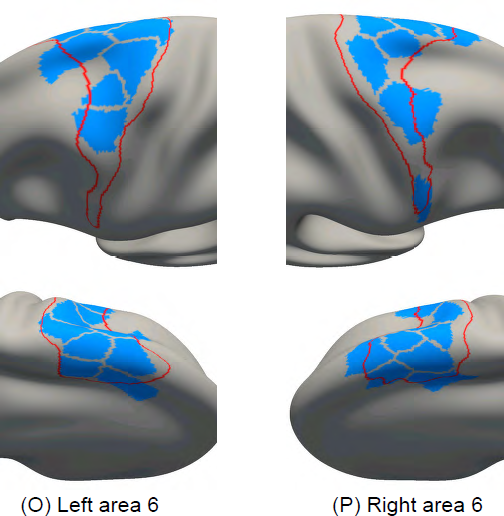
Parcels (blue) of the 400-area cerebral cortex parcellation overlaid on (red) boundaries of histologically defined areas

**Figure S10.**
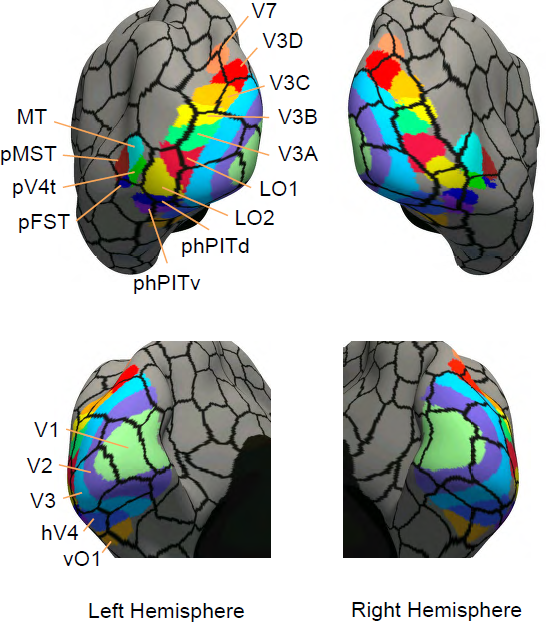
18 Visuotopic areas (Abdollahi et al., 2014) overlaid on (black) boundaries of the 400-area cerebral cortex parcellation.

**Figure S11A.**
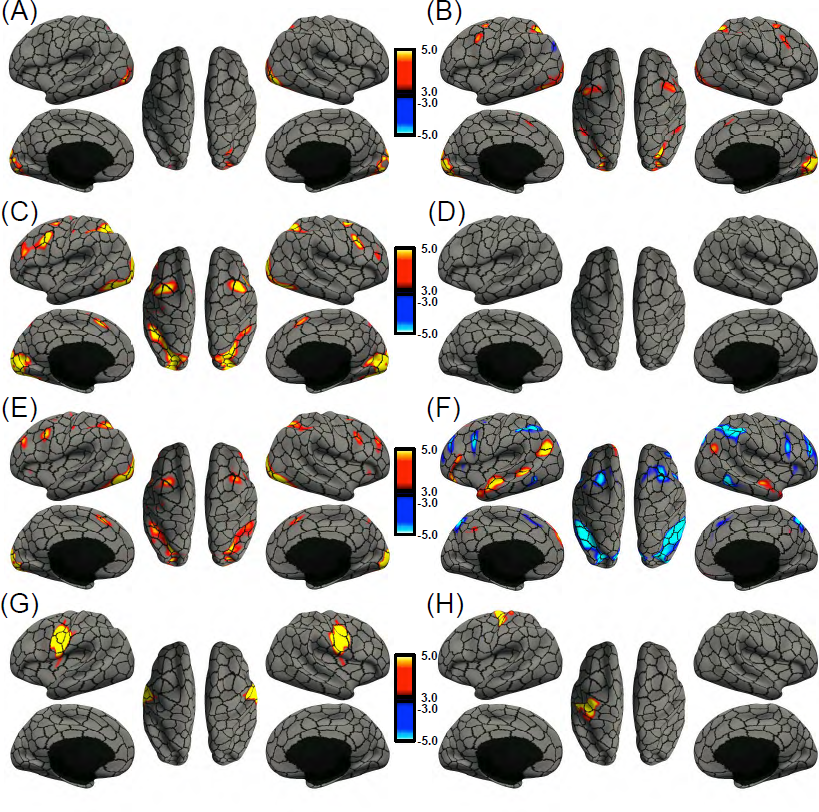
Group-average task activation maps of (A) emotion (faces – fixation), (B) gambling (punishment – fixation), (C) relational (matching – fixation), (D) social (theory of mind – random) and (E) working memory (2 back body – fixation), (F) language (story – math), (G) tongue motion (tongue – average motor), (H) right finger tapping (right finger – average motor) from the HCP dataset overlaid on (black) boundaries of 400-area cerebral cortex parcellation.

**Figure S11B.**
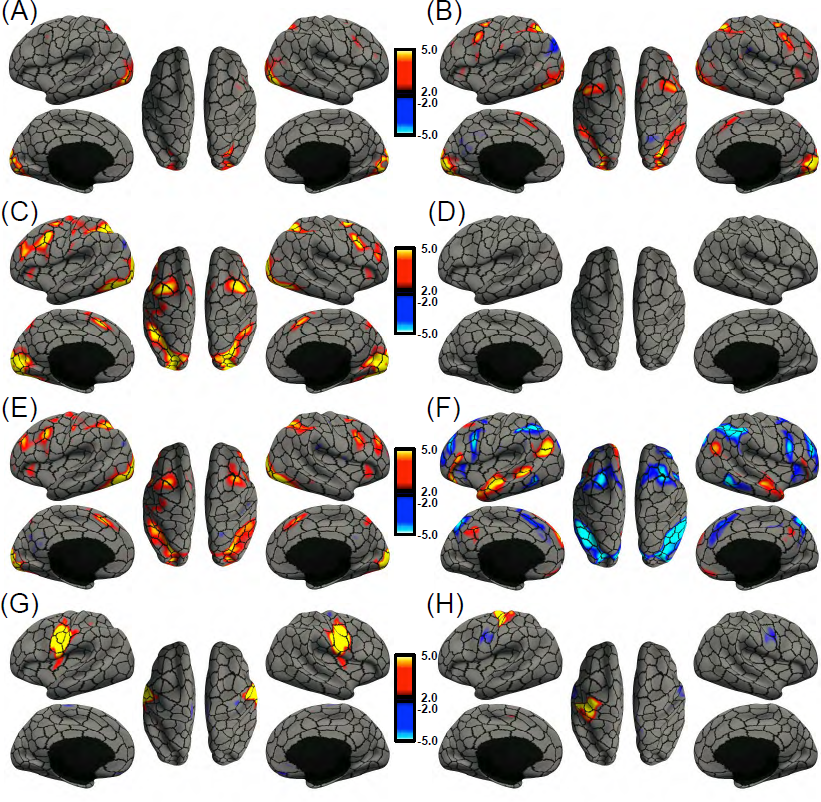
Group-average task activation maps of (A) emotion (faces – fixation), (B) gambling (punishment – fixation), (C) relational (matching – fixation), (D) social (theory of mind – random) and (E) working memory (2 back body – fixation), (F) language (story – math), (G) tongue motion (tongue – average motor), (H) right finger tapping (right finger – average motor) from the HCP dataset overlaid on (black) boundaries of 400-area cerebral cortex parcellation.

**Figure S11C.**
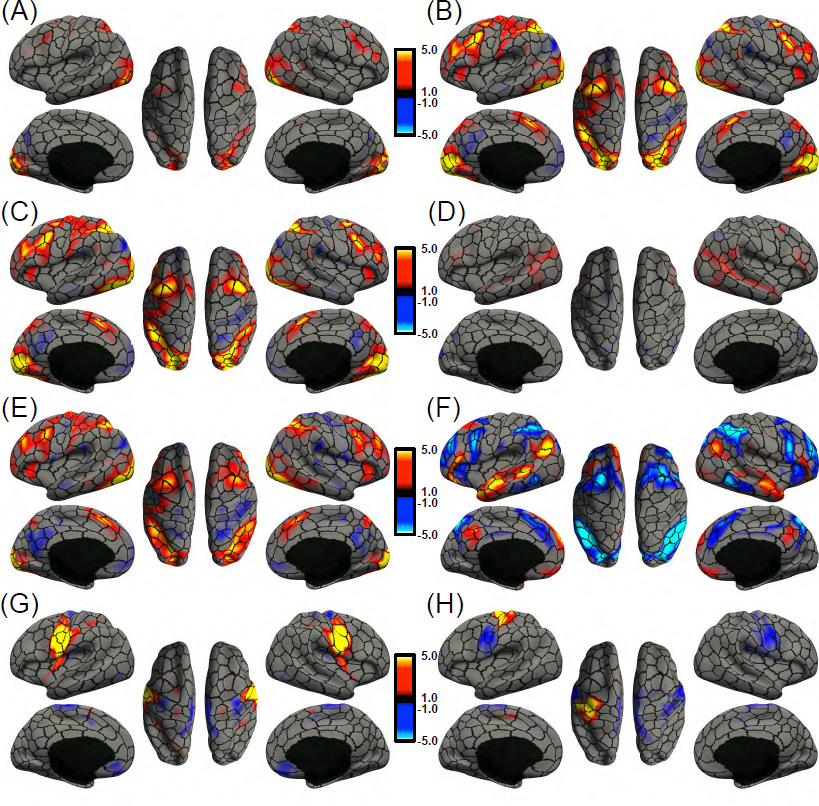
Group-average task activation maps of (A) emotion (faces – fixation), (B) gambling (punishment – fixation), (C) relational (matching – fixation), (D) social (theory of mind – random) and (E) working memory (2 back body – fixation), (F) language (story – math), (G) tongue motion (tongue – average motor), (H) right finger tapping (right finger – average motor) from the HCP dataset overlaid on (black) boundaries of 400-area cerebral cortex parcellation.

**Figure S11D.**
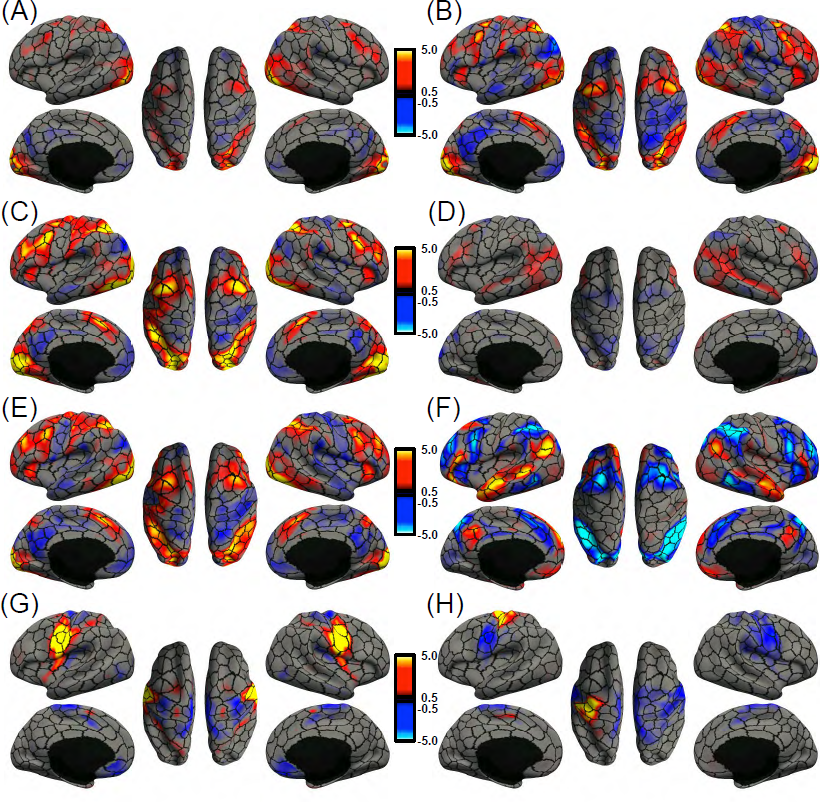
Group-average task activation maps of (A) emotion (faces – fixation), (B) gambling (punishment – fixation), (C) relational (matching – fixation), (D) social (theory of mind – random) and (E) working memory (2 back body – fixation), (F) language (story – math), (G) tongue motion (tongue – average motor), (H) right finger tapping (right finger – average motor) from the HCP dataset overlaid on (black) boundaries of 400-area cerebral cortex parcellation.

**Figure S11E.**
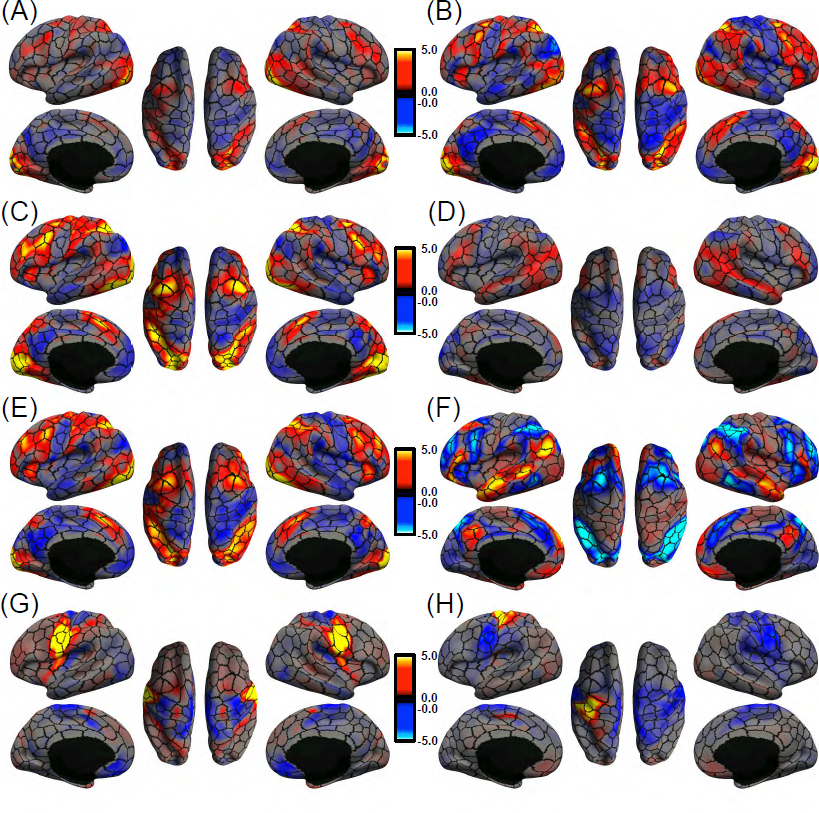
Group-average task activation maps of (A) emotion (faces – fixation), (B) gambling (punishment – fixation), (C) relational (matching – fixation), (D) social (theory of mind – random) and (E) working memory (2 back body – fixation), (F) language (story – math), (G) tongue motion (tongue – average motor), (H) right finger tapping (right finger – average motor) from the HCP dataset overlaid on (black) boundaries of 400-area cerebral cortex parcellation.

**Figure S12.**
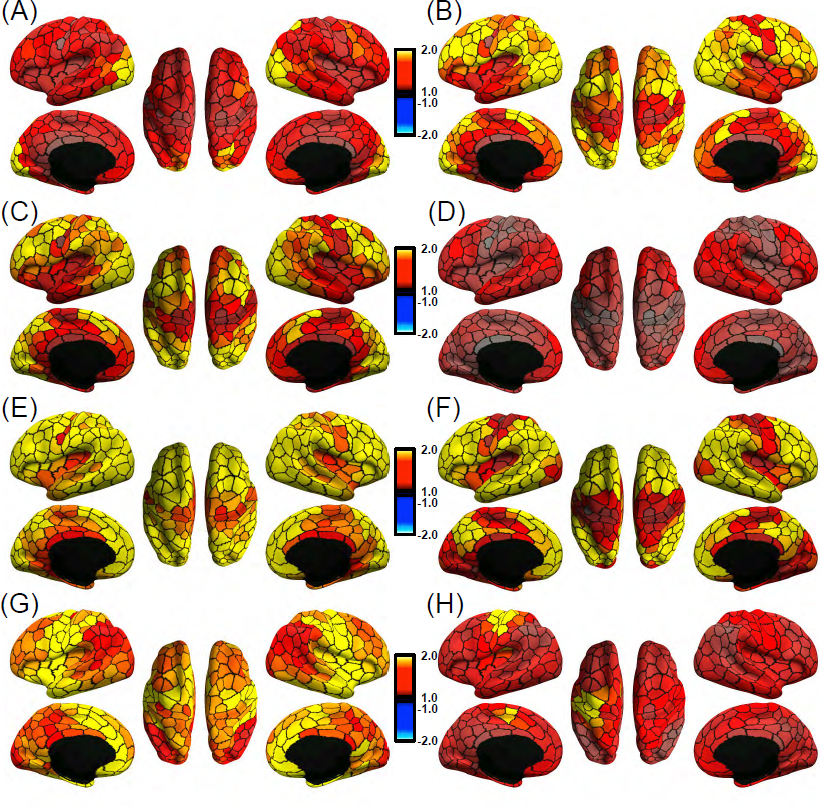
Functional inhomogeneity of each parcel (as measured by standard deviation of task activation z-scores) of the 400-area parcellation averaged across all HCP participants. (A) emotion (faces – fixation), (B) gambling (punishment – fixation), (C) relational (matching – fixation), (D) social (theory of mind – random) and (E) working memory (2 back body – fixation), (F) language (story – math), (G) tongue motion (tongue – average motor), (H) right finger tapping (right finger – average motor).

**Figure S13.**
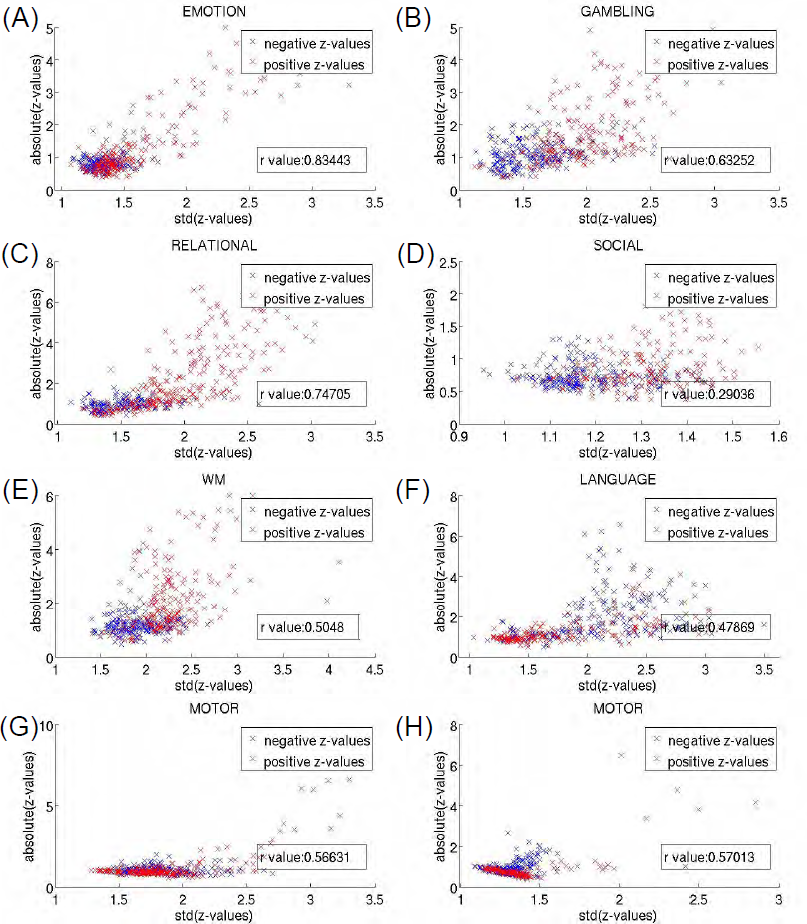
Scatterplot of each parcel’s task activation magnitude (absolute value of average z-scores) versus its functional inhomogeneity (as measured by standard deviation of z-scores) averaged across all HCP participants. Each cross in the scatterplot represents a parcel of the 400-area gwMRF parcellation. (A) emotion (faces – fixation), (B) gambling (punishment – fixation), (C) relational (matching – fixation), (D) social (theory of mind-random) and (E) working memory (2 back body – fixation), (F) language (story – math), (G) tongue motion (tongue – average motor), (H) right finger tapping (right finger – average motor). In general, stronger (positive or negative) activation is correlated with higher functional inhomogeneity.

**Figure S14.**
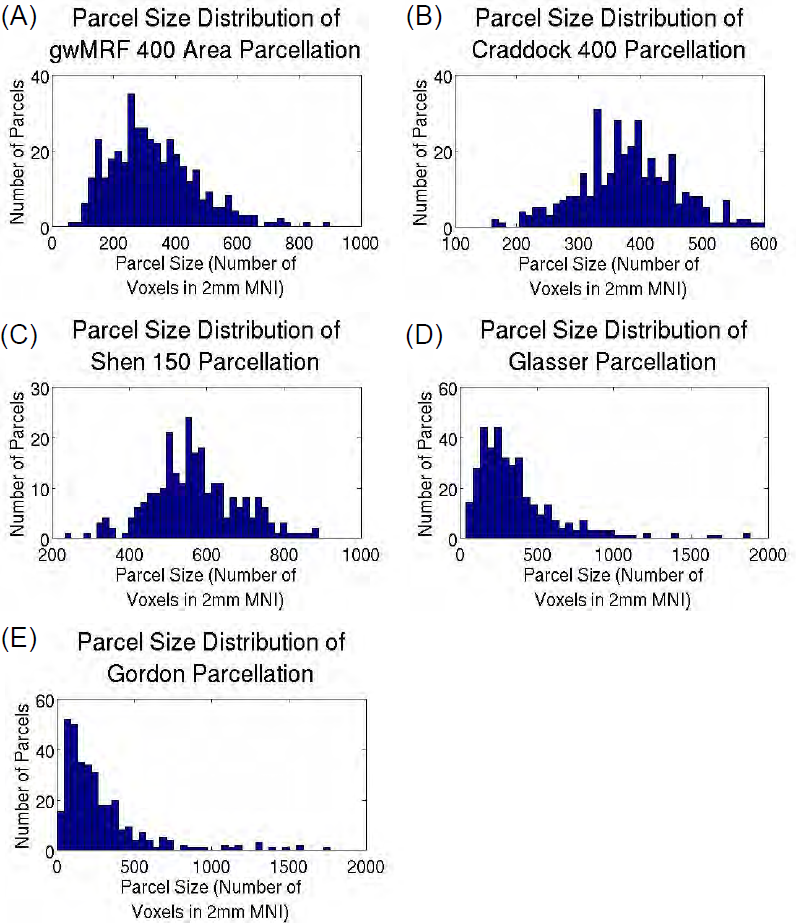
Parcel volume distributions for (A) gwMRF 400-area parcellation, (B) Craddock 400-area parcellation, (C) Shen 150-area parcellation, (D) Glasser 360-area parcellation and (E) Gordon 333-area parcellation. Volume of each parcel is computed as the number of voxels in 2mm MNI space. Ratio of largest and smallest parcels (A) 16 (B) 3.8 (C) 3.8 (D) 45 (E) 443

## Supplementary Material

This supplementary material is divided into Supplementary Methods and Supplementary Results to complement the Methods and Results sections in the main text, respectively.

## Supplementary Methods

This section provides additional mathematical and implementation details of the gwMRF parcellation procedure. Section S1 provides mathematical details about the gwMRF objective function. Section S2 summarizes the algorithm for optimizing the gwMRF objective function. Section S3 explains why time series normalization and concatenation is equivalent to Pearson’s correlation of time series for each subject followed by averaging across subjects. Section S4 provides derivation of the algorithm summarized in Section S2. Section S5 describes a computational trick to dramatically speed up the algorithm in Section S2, while decreasing memory usage. Section S6 provides details on how parameters of the gwMRF objective function are set.

### S1.Mathematical Model

In this section, we describe the gradient-weighted MRF (gwMRF) model for group-wise parcellation of the cerebral cortex. We assume a common surface coordinate system, where the cerebral cortex is represented by left and right hemisphere spherical meshes (i.e., FreeSurfer fsaverage surface meshes). Each mesh consists of a collection of vertices and edges connecting neighboring vertices into triangles (https://en.wikipedia.org/wiki/Triangle_mesh).

Let *q_n_* denote the *n*-th vertex, *N* denote the total number of vertices, and *N_qn_* denote the neighboring vertices of *q_n_* (as defined by the cortical meshes). Associated with each vertex *q_n_* is a 3×1 vector *s_n_* consisting of its 3-dimensional coordinates on the spherical mesh and preprocessed fMRI time course for each participant.

The fMRI time course of each participant at each vertex was normalized to be of zero mean and standard deviation of one. For participants with two runs, each run was normalized separately. For each vertex *q_n_*, the normalized time courses of all runs of all subjects were concatenated into a long (column) vector and normalized to unit norm. The resulting normalized time course is denoted as *y_n_*. This normalization procedure is motivated by the fact that the resulting inner product between two time courses *y_n_* and *y_m_* is equivalent to computing Pearson product-moment correlation coefficient for each participant and then averaging across the participants (more details in Supplementary Methods S3).

To summarize, the input data consisted of normalized fMRI time courses {*y_1_,…, y_w_*} denoted as *y_1:N_* and 3-dimensional spherical coordinates *s*_1:*N*_. Our goal is to estimate a population-level parcellation label *l_n_* at each vertex *q_n_*, where *l_n_* ∈ {1,…, *L*}. The complete parcellation {*l_1_,…, l_N_*} is denoted as *l*_1:*N*_. The following gwMRF model specifies the joint probability distribution of labels *l*_1:N_, time courses *y*_1:*N*_ and spatial positions *s*_1:*N*_:

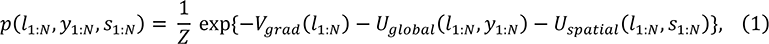

where *Z* is a normalization term to ensure *p*(*l*_1:*N*_, *y*_1:*N*_, *s*_1:*N*_) is a valid probability distribution. *V_grad_ l*_1:*N*_) consists of pairwise potentials incorporating the local gradient approach, *U_global_*(*l*_1:*N*_, *y*_1:*N*_) consists of unary potentials encoding the global similarity approach and *U_spatial_* (*l*_1:*N*_, *S*_1:*N*_) consists of unary potentials ensuring parcels remain spatially connected.

More specifically, *V_grad_*(*l*_1:*N*_) penalizes neighboring vertices with different labels, but the penalties are weighted so that there is lower penalty in the presence of local gradients in RSFC (hence the name gradient weighted MRF):

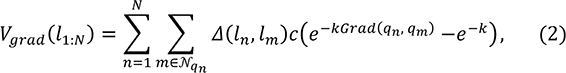

where Δ(*l_n_, l_m_*) is equal to one if *l_n_* ≠ *l_m_* and zero otherwise. *Grad(q_n_, q_m_*) is the RSFC gradient magnitude between vertices *q_n_* and *q_m_*, with higher values indicating stronger gradients. In this work, we utilize the state-of-the-art gradients computed by Gordon et al. (2016), where *Grad(q_n_, q_m_)* ranges from zero to one. *k* and *c* are tunable parameters both greater than zero. If *l_n_ = l_m_*, the penalty is always zero. If *l_n_* ≠ *l_m_* and *Grad(q_n_, q_m_*) = 0 (i.e., zero gradient), then the penalty is *c*(1 – *e^-k^*). As *Grad(q_n_, q_m_*) increases from zero to one, the penalty decays exponentially to zero. Therefore *k* controls the exponential decay rate, while *c* controls the overall magnitude of the penalty and therefore the weight of the local gradients relative to the other terms in the MRF.

The global connectivity similarity approach is encoded by *U_Giobal_*(*l*_1:*n*_, *y*_1:*N*_) defined as

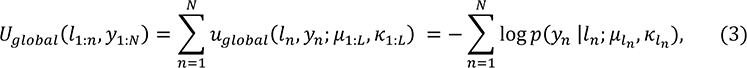

where *p(y_n_|l_n_;μl_n_,kl_n_)* follows a von Mises-Fisher distribution:

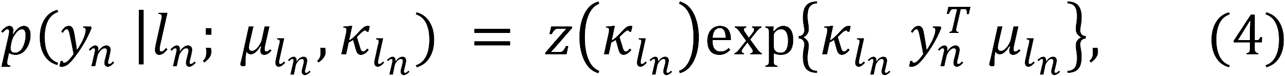

and *μ_l_n__ and K_l_n__* are the mean direction and concentration parameter of the *l_n_*-th von Mises-Fisher distribution and *z*(*K_l_n__*) is a normalization constant. As will be made clear (see Supplementary Methods S2), we can think of *μ_l_* as the mean time course of parcel *l*, normalized to unit norm. Therefore if *y_n_* is similar to μ_l_(i.e., 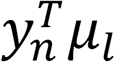 is big), then vertex *q_n_* is more likely to be assigned to parcel *l. U_global_*(*l*_1:*n*_, *y*_1:*N*_) is the analogue of the von Mises-Fisher mixture model employed in Yeo et al. (Yeo et al. 2011), and thus encodes the global connectivity similarity approach.

With just *V_grad_* and *U_global_*, many parcels will be spatially distributed due to strong long-range RSFC. Requiring parcels in a MRF framework to be spatially connected (with minimal other assumptions) is non-trivial (Nowozin and Lampert 2010; Honnorat et al. 2015). Here, spatial connectedness is imposed by *U_spatial_* defined as

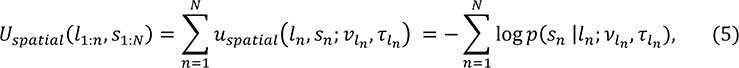

where *p(s_n_ | l_n_; v_l_n__, τ_l_n__)* follows a von Mises-Fisher distribution:

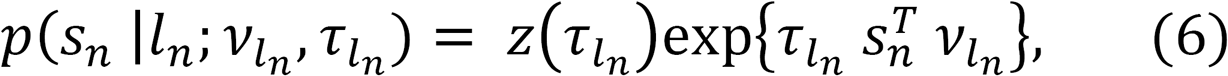

where *v__l_n_* is the mean spatial direction of parcel *l_n_* and τ_1:L_ are tunable parameters greater than zero. As will be made clear (see Supplementary Methods S2), we can think of *v_l_* as the mean spatial coordinates of parcel *l* normalized to unit norm (i.e., sphere). Therefore if vertex *q_n_* is spatially close to the mean spatial location of parcel *l* (i.e., 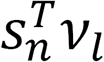 is big), then vertex *q_n_* is more likely to be assigned to parcel *l.* Consequently, for large enough values of τ_1:*L*_, the parcels will be spatially connected. This approach is significantly less computationally expensive than competing approaches (Nowozin and Lampert 2010; Honnorat et al. 2015). However, a serious problem is that large values of τ_1:*L*_ lead to round parcels, which are not biologically realistic. For example, we expect cortical areas in the cingulate to be long and narrow (Vogt 2009). To avoid this issue, the optimization procedure (see Supplementary Methods S2) starts with large values of τ_1:*L*_, and then iteratively decreases τ_1:*L*_, thus ensuring spatially connected parcels that are not constrained to be round.

### S2.Model Inference

Given observed time courses *y*_1:*N*_, spatial positions *S*_1:*N*_ and for a fixed set of parameters *c, k* and τ_1:*L*_, we seek to estimate {*l*_1:*N*_, μ_1:*L*_, *K*_1:*L*_, *v*_1:*L*_} using the maximum-a-posteriori (MAP) principle:

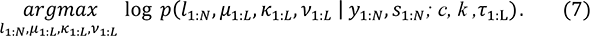

The optimization is achieved by coordinate descent. At each iteration, we use the current estimates of {*μ*_1:*L*_, *k*_1:*L*_, *v*_1:*L*_) to estimate labels *l*_1:*N*_ using graph cuts (Delong et al. 2010). We then use the estimates of labels *l*_1:*N*_ to infer {*μ*_1:*L*_, *k*_1:*L*_, *v*_1:*L*_):

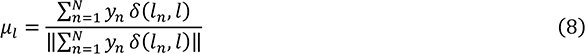

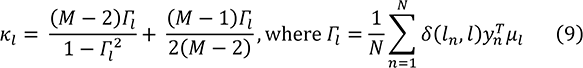

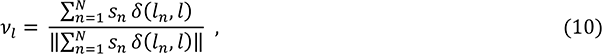

where *δ*(*l_n_,l)* is one if *l_n_ = l,* and is zero otherwise, ||⋅|| corresponds to the *l*_2_-norm and *M* is the length of *y_n_*. Therefore the estimate of *μ_l_* is the average time course of vertices constituting parcel *l,* normalized to be unit norm. Similarly, the estimate of *v_l_* is the average spatial locations of vertices constituting parcel *l,* normalized to be unit norm. Detailed derivations are provided in Supplementary Methods S4. The algorithm proceeds by initializing {*μ*_1:*L*_, *K*_1:*L*_, *V*_1:*L*_} and then iterating graph cuts (to estimate *l*_1:*N*_) and Eqs. (8) to (10) until convergence. Supplementary Methods S5 describes a trick to reduce memory requirements of the algorithm and speed it up by almost two orders of magnitude. For future reference, we refer to this algorithm as the **MAPI** algorithm.

As discussed in the previous section, large values of τ_1:*L*_ lead to spatially connected parcels that are unrealistically round. To resolve this issue, for a fixed set of parameters *c* and *k,* we first set τ_1:*L*_ to a sufficiently large constant to ensure parcels are spatially connected. After the MAP1 algorithm converges, we repeatedly divide the τ of each parcel by five and re-run the MAP1 algorithm using the previous estimate of {*μ*_1:*L*_, *k*_1:*L*_, *v*_1:*L*_) as initialization. If decreasing τ causes some parcels to become spatially distributed, we repeatedly quintuple the τ of each spatially distributed parcel (while keeping the τ of each spatially connected parcel constant) and re-run the MAP1 algorithm until all parcels are again spatially connected. The whole process of reducing and doubling τ is repeated until τ_1:*L*_ = 0 or we detect a repeated setting of τ_1:*L*_. For future reference, we refer to this algorithm as the **MAP2** algorithm. While there is no theoretical guarantee, we find that the τ’s of almost all parcels are driven to zero in practice. Visual inspection also suggests that the resulting parcels are not constrained to be round.

Finally, for a fixed set of parameters *c* and *k,* and a large initial setting of τ_1:*L*_, we randomly initialize {*μ*_1:*L*_, *K*_1:*L*_, *V*_1:*L*_), and run the MAP2 algorithm until convergence. This is repeated with multiple random initializations of {*μ*_1:*L*_, *K*_1:*L*_, *V*_1:*L*_}. Random initialization is achieved by setting *μ*_1:L_ and *v*_1:*L*_ to be the time courses and spatial locations of *L* randomly selected vertices respectively. *k*_1:*L*_ is initialized to be 12,500. The parcellation with the highest likelihood (Eq. (1) excluding *U_spatial_*) is selected as the final solution. For future reference, we refer to this algorithm as the MAP3 algorithm. The **MAP3** algorithm is utilized in all subsequent experiments and evaluation.

In this work, the parcellation is performed separately for the left and right hemispheres. The number of labels *L* is already discussed in the *Methods* section of the main text. The setting of parameters *c* and *k,* initial setting of τ_1:*L*_, and the number of random initializations are discussed in Supplementary Methods S6.

### Data Normalization

In this work, the fMRI time course of each participant at each vertex is normalized with mean of zero and standard deviation of one. For each vertex, the normalized time courses of all runs of all subjects are concatenated into a long (column) vector and normalized to unit norm. Assuming the fMRI data of each participant is of length *T,* then this section demonstrates that the inner product between two normalized time courses is the same as computing Pearson product-moment correlation coefficient for each participant and then averaging across the participants.

Let *a_st_* and *b_st_* be the fMRI signals of subject *s* at time *t* at two vertices. The Pearson product-moment correlation coefficient between the two time courses is given by

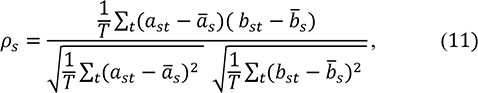

where 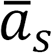 and 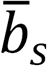 are the mean of *a_st_* and *b_st_* respectively (for each subject *s*). Suppose *a_st_* and *b_st_* have been normalized to mean zero and standard deviation one (for each subject *s*), then Eq. (11) becomes

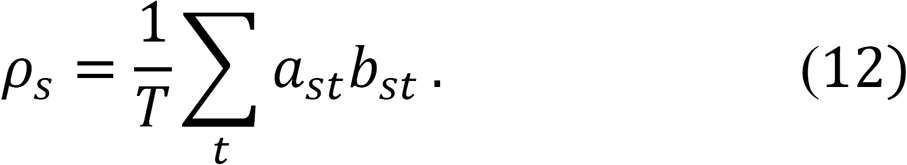

The mean correlation across participants is given by

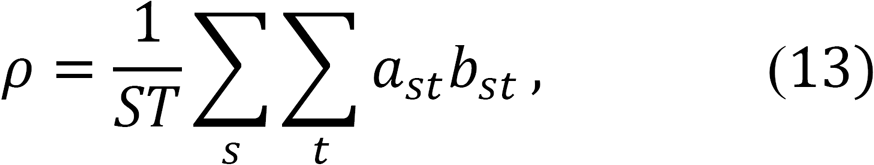

which can be written as

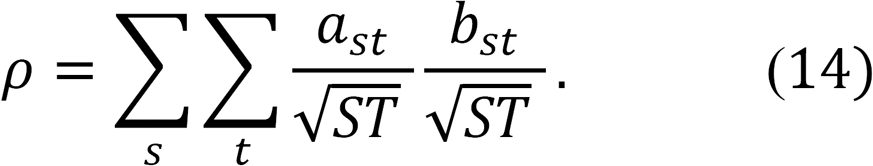

If we concatenate *a_st_* and *b_st_* across all subjects and timepoints, and denote the resulting time courses as *e_t_* and *f_t_* respectively, then Eq. (14) is equivalent to

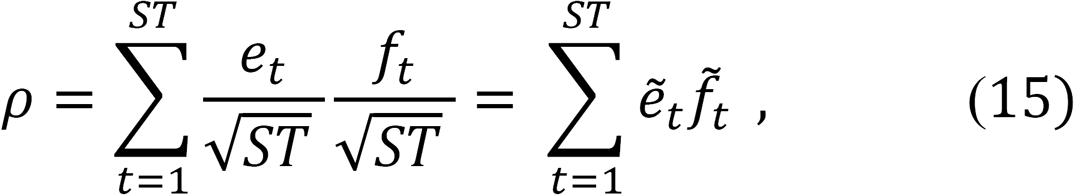

where 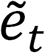 and 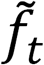 are unit norm signals obtained by dividing *e_t_* and *f_t_* by 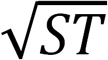. To see this, recall that *a_st_* and *b_st_* have been normalized to mean zero and standard deviation one (for each subject *s*). Therefore the *l_2_*-norm of *a_st_* and *b_st_* (for each subject *s*) is equal to 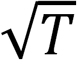. This means that the *l_2_*-norm of the concatenated time courses *e_t_* and *f_t_* is equal to 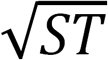. Dividing *e_t_* and *f_t_* by 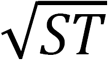 results in unit norm signals 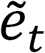 and 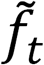. Therefore the mean correlation across subjects *p* is equal to the inner product of the unit norm time courses 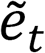 and 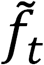 (Eq. (15)).

In this work, the GSP dataset consisted of subjects with one or two runs. Each run is separately normalized to a mean of zero and a standard deviation of one. Mathematically, this means that subjects with two runs are weighted more than subjects with one run, which might make sense given that subjects with more data should enjoy better signal to noise ratio. Given the large dataset (N = 1,489), we think it is unlikely that the final parcellation will reflect the idiosyncrasies of any particular subject. Furthermore, the evaluation metrics were computed for each subject and then averaged. Therefore any concerns about the uneven weighting would not affect the evaluation.

### MAP Estimation

This section derives the MAP1 algorithm (Supplementary Methods S2). Recall that given observed time courses *y*_1:*N*_, spatial positions *S*_1:*N*_ and for a fixed set of parameters *c, k* and *t*_1:*L*_, we seek to estimate *l*_1:*N*_, *μ*_1:*L*_, *k*_1:*l*_, *v*_1:*L*_ using maximum-a-posteriori (MAP) principle as stated in Eq. (7) in Supplementary Methods S2:

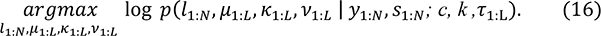

Assuming a uniform prior on *μ*_1:*L*_, *k*_1:*l*_, *v*_1:*L*_ we can write Eq. (16) as

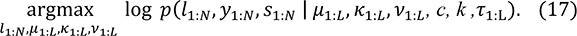

We can then plug in Eq. (1) and write

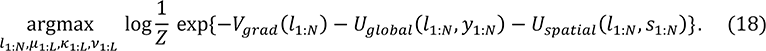

The partition function Z is a function of the fixed parameters *c, k* and τ_1:*L*_, and so can be dropped from the optimization defined in Eq. (18). In addition, by plugging Eqs. (2) to (6) into Eq. (18), we get

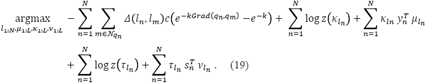

The global optimum of Eq. (19) is NP-hard to estimate. We optimize Eq. (19) by coordinate descent, which guarantees convergence to a local optimum.

At each iteration, we fix the current estimates of {*μ*_1:*L*_, *k*_1:*L*_, *v*_1:*L*_}, and optimize Eq. (19) to estimate labels *l*_1:*N*_. By flipping the sign of Eq. (19), the maximization problem becomes a minimization problem:

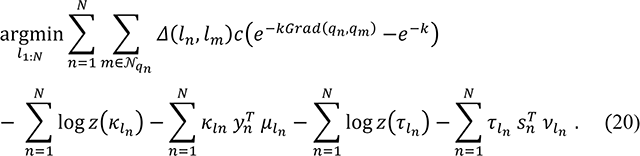

The pairwise potentials in Eq. (20) are submodular. However, the unary potentials are negative, so graph cuts (Delong et al. 2010) cannot be directly applied because the resulting max-cut sub-problem is intractable since the graph has both positive and negative weights. Therefore to apply graph cuts, we add a positive constant *C* to all the unary potentials so all the weights become positive. Additionally, we divide by the number of vertices *N* to avoid a numerical overflow:

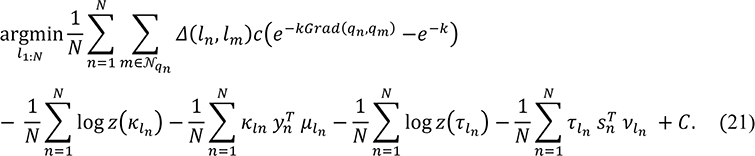

Eq. (21) can be directly optimized using graph cuts (Delong et al. 2010).

We then fix the current estimates of labels *l*_1:*N*_ and optimize Eq. (19) to estimate {*μ*_1:*L*_ *K*_1:*L*_, *v*_1:*L*_). For fixed label estimates *l*_VN_, optimization of Eq. (19) for {*μ*_1:*L*_, *k*_1:*L*_, *v*_1:*L*_} is equivalent to the maximum likelihood estimation of the von Mises-Fisher distributions. By using the constraints that 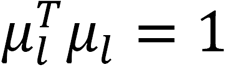, 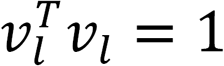 and *K* > 0, and differentiating Eq. (19) with respect to *μ_l_, K_l_* and *V_l_*, and setting the derivatives to zero, we get the following update equations (Lashkari et al. 2010):

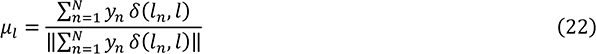

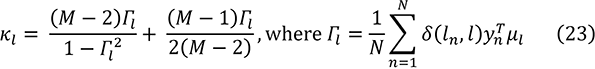

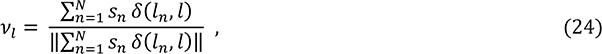

where *δ*(*l_n_, l*) is one if *l_n_ = l,* and is zero otherwise, and ||⋅|| corresponds to the *l_2_*-norm. Therefore the MAP1 algorithm proceeds by initializing {*μ*_1:*L*_, *k*_1:*L*_, *v*_1:*L*_} and then iterating between Eq. (21) and Eqs. (22) to (24) until convergence.

### S5. Speeding Up the MAP1 Algorithm

This section describes an implementation trick that speeds up the MAP1 algorithm by a factor of approximately 80 for the full GSP dataset. The resulting implementation enjoys computational and memory complexities independent of the number of fMRI time points and the number of subjects. Recall that the fMRI data consists of normalized time course *y_n_* at each vertex *q_n_.* By utilizing Eq. (22), observe that

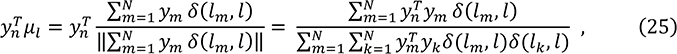

implying that the inner product between time course *y_n_* and mean direction *μ_l_* can be computed with just the inner products of the input time courses.

Recall that the MAP1 algorithm iterates between Eq. (21) and Eqs. (22) to (24) until convergence. The fMRI time courses {*y_x_,…, y_N_*} and mean cluster directions {*μ_l_,…, μ_L_*} appear in Eq. (21) and Eq. (23) only in the form 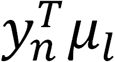 for different values of *n* and *l.* Eq. (25) implies that Eq. (21) and Eqs. (23) can be computed with just the inner products of the fMRI time courses without needing the actual time courses themselves. Therefore our actual implementation of the MAP1 algorithm iterates between Eq. (21), Eq. (23) and Eq. (24), without explicitly computing Eq. 22). Instead of passing in the normalized fMRI time courses, representable by a matrix *Y* = [*y_1_*…*y_N_*] of dimensions *T* × *N*, the inner product matrix *A* = *Y^T^Y* was precomputed and passed in as an input to the algorithm. The matrix *A* is of size *N* × *N,* so we achieved substantial savings because *N =* 40962 for each fsaverage6 spherical mesh and *T* is 308,640 for the full GSP dataset.

### S6. Implementation Details of the gwMRF Model

There are four “free” parameters in the gwMRF model. The number of parcels *L* is already discussed in the *Methods* section of the main text. This section provides more details on how the local gradient weight *c* (Eq. (2)), exponential decay parameter *k* (Eq. (2)) and the initial weight of the spatial connectedness term τ_1:*L*_ (Eq. (4)) are set.

For comparison with other publicly available parcellations, the MAP3 algorithm (Supplementary Methods S2) was applied to the GSP training set (N = 744). The number of parcels *L* was set to match the number of parcels in the publicly available parcellation. We explored the local gradient weight *c* from 25,000 to 500,000 and the exponential decay parameter *k* from 1 to 30. Based on visual inspection of the parcellations and functional connectivity homogeneity in the GSP training set, the gradient weight *c* and exponential decay parameter *k* were set to 1e5 and 15 respectively for all comparisons. The initial weight of the spatial connectedness term τ_1:L_ was set to be 5e6. The MAP3 algorithm was run with 500 random initializations.

Parcellations at multiple resolutions (400, 600, 800 and 1,000 areas) were generated from the full GSP dataset for public distribution, as well as from the GSP training and test sets for stability analyses. With the change in the number of parcels, the relative weights of the various terms in the gwMRF model (Eq. (1)) needed to be modified. Therefore we could not use exactly the same parameters as before. Based on visual inspection of the resulting parcellations, the gradient weight *c* was set to 150,000 for the 400-area parcellation and 300,000 for 600 or more parcels. The exponential decay parameter *k* was set to 15 for 400-area parcellation and 7.5 for 600 or more parcels. The initial weight of the spatial connectedness term τ_1:L_ was set to be 5e8. MAP3 algorithm was run with 1000 random initializations.

## References

Abdollahi RO, Kolster H, Glasser MF, Robinson EC, Coalson TS, Dierker D, Jenkinson M, Van Essen DC, Orban GA. 2014. Correspondences between retinotopic areas and myelin maps in human visual cortex. Neuroimage. 99:509–524.

Amunts K, Malikovic A, Mohlberg H, Schormann T, Zilles K. 2000. Brodmann’s areas 17 and 18 brought into stereotaxic space—where and how variable? Neuroimage. 11:66–84.

Amunts K, Schleicher A, Buergel U, Mohlberg H, Uylings HBM, Zilles K. 1999. Broca’s region revisited: Cytoarchitecture and intersubject variability. J Comp Neurol. 412:319–341.

Amunts K, Weiss PH, Mohlberg H, Pieperhoff P, Eickhoff S, Gurd JM, Marshall JC, Shah NJ, Fink GR, Zilles K. 2004. Analysis of neural mechanisms underlying verbal fluency in cytoarchitectonically defined stereotaxic space—the roles of Brodmann areas 44 and 45. Neuroimage. 22:42–56.

Amunts K, Zilles K. 2015. Architectonic mapping of the human brain beyond Brodmann. Neuron. 88:1086–1107.

Arslan S, Ktena SI, Makropoulous, Robinson EC, Rueckert D, Parisot S. 2017. Human brain mapping: a systematic comparison of parcellation methods for the human cerebral cortex. In press

Baker JT, Holmes AJ, Masters GA, Yeo BTT, Krienen F, Buckner RL, Öngür D. 2014. Disruption of cortical association networks in schizophrenia and psychotic bipolar disorder. JAMA psychiatry. 71:109–118.

Barch DM, Burgess GC, Harms MP, Petersen SE, Schlaggar BL, Corbetta M, Glasser MF, Curtiss S, Dixit S, Feldt C, Nolan D, Bryant E, Hartley T, Footer O, Bjork JM, Poldrack R, Smith S, Johansen-Berg H, Snyder AZ, Van Essen DC. 2013. Function in the human connectome: Task-fMRI and individual differences in behavior. Neuroimage. 80:169–189.

Barrett LF, Satpute AB. 2013. Large-scale brain networks in affective and social neuroscience: towards an integrative functional architecture of the brain. Curr Opin Neurobiol. 23:361–372.

Beckmann CF, DeLuca M, Devlin JT, Smith SM. 2005. Investigations into resting-state connectivity using independent component analysis. Philos Trans R Soc B Biol Sci. 360:1001–1013.

Behzadi Y, Restom K, Liau J, Liu TT. 2007. A component based noise correction method (CompCor) for BOLD and perfusion based fMRI. Neuroimage. 37:90–101.

Bellec P, Rosa-Neto P, Lyttelton OC, Benali H, Evans AC. 2010. Multi-level bootstrap analysis of stable clusters in resting-state fMRI. Neuroimage. 51:1126–1139.

Bertolero MA, Yeo BTT, D’Esposito M. 2015. The modular and integrative functional architecture of the human brain. Proc Natl Acad Sci. 112:E6798–E6807.

Betzel RF, Avena-Koenigsberger A, Goñi J, He Y, De Reus MA, Griffa A, Vértes PE, Mišic B, Thiran J-P, Hagmann P, others. 2016. Generative models of the human connectome. Neuroimage. 124:1054–1064.

Betzel RF, Byrge L, He Y, Goñi J, Zuo X-N, Sporns O. 2014. Changes in structural and functional connectivity among resting-state networks across the human lifespan. Neuroimage. 102:345–357.

Biswal B, Zerrin Yetkin F, Haughton VM, Hyde JS. 1995. Functional Connectivity in the Motor Cortex of Resting Human Brain Using Echo-Planar MRI. Magn Reson Med. 34:537–541.

Blumensath T, Jbabdi S, Glasser MF,Van Essen DC, Ugurbil K, Behrens TEJ, Smith SM. 2013. Spatially constrained hierarchical parcellation of the brain with resting-state fMRI. Neuroimage. 76:313–324.

Brodmann K. 1909. Vergleichende Lokalisationslehre der Grosshirnrinde: in ihren Prinzipien dargestellt auf Grund des Zellenbaues. Ja Barth.

Buckner RL, Krienen FM. 2013. The evolution of distributed association networks in the human brain. Trends Cogn Sci. 17:648–665.

Buckner RL, Krienen FM, Castellanos A, Diaz JC, Yeo BTT. 2011. The organization of the human cerebellum estimated by intrinsic functional connectivity. J Neurophysiol. 106:2322–2345.

Buckner RL, Krienen FM, Yeo BTT. 2013. Opportunities and limitations of intrinsic functional connectivity MRI. Nat Neurosci. 16:832–837.

Buckner RL, Yeo BTT. 2014. Borders, map clusters, and supra-areal organization in visual cortex. Neuroimage. 93:292–297.

Bullmore E, Sporns O. 2009. Complex brain networks: graph theoretical analysis of structural and functional systems. Nat Rev Neurosci. 10:186–198.

Bzdok D, Laird AR, Zilles K, Fox PT, Eickhoff SB. 2013. An investigation of the structural, connectional, and functional subspecialization in the human amygdala. Hum Brain Mapp. 34:3247–3266.

Calhoun VD, Kiehl KA, Pearlson GD. 2008. Modulation of temporally coherent brain networks estimated using ICA at rest and during cognitive tasks. Hum Brain Mapp. 29:828–838.

Cao M, Wang J-H, Dai Z-J, Cao X-Y, Jiang L-L, Fan F-M, Song X-W, Xia M-R, Shu N, Dong Q, others. 2014. Topological organization of the human brain functional connectome across the lifespan. Dev Cogn Neurosci. 7:76–93.

Choi EY, Yeo BTT, Buckner RL. 2012. The organization of the human striatum estimated by intrinsic functional connectivity. J Neurophysiol. 108: 2242–2263.

Churchland PS, Sejnowski TJ. 1988. Perspectives on cognitive neuroscience. Science. 242:741–745.

Cohen AL, Fair D a, Dosenbach NUF, Miezin FM, Dierker D, Van Essen DC, Schlaggar BL, Petersen SE. 2008. Defining functional areas in individual human brains using resting functional connectivity MRI. Neuroimage. 41:45–57.

Cole MW, Bassett DS, Power JD, Braver TS, Petersen SE. 2014. Intrinsic and task-evoked network architectures of the human brain. Neuron. 83:238–251.

Corbetta M, Shulman GL. 2002. Control of goal-directed and stimulus-driven attention in the brain. Nat Rev Neurosci. 3:201–15.

Craddock RC, James GA, Holtzheimer PE, Hu XP, Mayberg HS. 2012. A whole brain fMRI atlas generated via spatially constrained spectral clustering. Hum Brain Mapp. 33:1914–1928.

Cragg BG. The topography of the afferent projections in the circumstriate visual cortex of the monkey studied by the Nauta method. Vision Res 9:733–747, 1969.

Damoiseaux JS, Rombouts SARB, Barkhof F, Scheltens P, Stam CJ, Smith SM, Beckmann CF. 2006. Consistent resting-state networks across healthy subjects. Proc Natl Acad Sci U S A. 103:13848–13853.

Davis C, Kleinman JT, Newhart M, Gingis L, Pawlak M, Hillis AE. 2008. Speech and language functions that require a functioning Broca’s area. Brain Lang. 105:50–58.

De Luca M, Beckmann CF, De Stefano N, Matthews PM, Smith SM. 2006. fMRI resting state networks define distinct modes of long-distance interactions in the human brain. Neuroimage. 29:1359–1367.

Deco G, Ponce-Alvarez A, Mantini D, Romani GL, Hagmann P, Corbetta M. 2013. Resting-state functional connectivity emerges from structurally and dynamically shaped slow linear fluctuations. J Neurosci. 33:11239–11252.

Delong A, Osokin A, Isack HN, Boykov Y. 2010. Fast approximate energy minimization with label costs. In: 2010 IEEE Computer Society Conference on Computer Vision and Pattern Recognition. IEEE. p. 2173–2180.

Ding S, Royall JJ, Sunkin SM, Ng L, Facer BAC, Lesnar P, Guillozet-Bongaarts A, McMurray B, Szafer A, Dolbeare TA, Stevens A, Tirrell L, Benner T, Caldejon S, Dalley RA, Dee N, Lau C, Nyhus J, Reding M, Riley ZL, Sandman D, Shen E, van der Kouwe A, Varjabedian A, Write M, Zollei l, Dang C, Knowles JA, Koch C, Phillips JW, Sestan N, Wohnoutka P, Zielke HR, Hohmann JG, Jones AR, Bernard A, Hawrylycz MJ, Hof PR, Fischl B, Lein ES. 2016. Comprehensive cellular-resolution atlas of the adult human brain: Adult human brain atlas. J Comp Neurol. 524:3127–3481.

Di Martino A, Scheres A, Margulies DS, Kelly AMC, Uddin LQ, Shehzad Z, Biswal B, Walters JR, Castellanos FX, Milham MP. 2008. Functional connectivity of human striatum: a resting state FMRI study. Cereb Cortex. 18:2735–2747.

Dobromyslin VI, Salat DH, Fortier CB, Leritz EC, Beckmann CF, Milberg WP, McGlinchey RE. 2012. Distinct functional networks within the cerebellum and their relation to cortical systems assessed with independent component analysis. Neuroimage. 60:2073–85.

Dosenbach NUF, Fair DA, Miezin FM, Cohen AL, Wenger KK, Dosenbach RAT, Fox MD, Snyder AZ, Vincent JL, Raichle ME, others. 2007. Distinct brain networks for adaptive and stable task control in humans. Proc Natl Acad Sci. 104:11073–11078.

Eickhoff SB, Stephan KE, Mohlberg H, Grefkes C, Fink GR, Amunts K, Zilles KA. 2005. A new SPM toolbox for combining probabilistic cytoarchitectonic maps and functional imaging data. NeuroImage. 1:1325–1335.

Eickhoff SB, Bzdok D, Laird AR, Roski C, Caspers S, Zilles K, Fox PT. 2011. Co-activation patterns distinguish cortical modules, their connectivity and functional differentiation. Neuroimage. 57:938–949.

Eickhoff SB, Constable RT, Yeo BTT. 2017. Topographic organization of the cerebral cortex and brain cartography. NeuroImage. In press

Eickhoff SB, Grefkes C. 2011. Approaches for the integrated analysis of structure, function and connectivity of the human brain. Clin EEG Neurosci. 42:107–121.

Eickhoff SB, Thirion B, Varoquaux G, Bzdok D. 2015. Connectivity-based parcellation: Critique and implications. Hum Brain Mapp. Epub ahead.

Fair D, Nigg JT, Iyer S, Bathula D, Mills KL, Dosenbach NUF, Schlaggar BL, Mennes M, Gutman D, Bangaru S, Buitelaar JK, Dickstein DP, Di Martino A, Kennedy DN, Kelly C, Luna B, Schweitzer JB, Velanova K, Wang Y-F, Mostofsky SH, Castellanos FX, Milham MP. 2013. Distinct Neural Signatures Detected for ADHD Subtypes After Controlling for Micro-Movements in Resting State Functional Connectivity MRI Data. Front Syst Neurosci. 6.

Fan L, Li H, Zhuo J, Zhang Y, Wang J, Chen L, Yang Z, Chu C, Xie S, Laird AR, others. 2016. The Human Brainnetome Atlas: A New Brain Atlas Based on Connectional Architecture. Cereb Cortex. 26:3508–3526.

Felleman DJ, Van Essen DC. 1991. Distributed hierarchical processing in the primate cerebral cortex. Cereb Cortex. 1:1–47.

Finn ES, Shen X, Scheinost D, Rosenberg MD, Huang J, Chun MM, Papademetris X, Constable RT. 2015. Functional connectome fingerprinting: identifying individuals using patterns of brain connectivity. Nat Neurosci. 18:1664–1671

Fischl B. 2012. FreeSurfer. Neuroimage. 62:774–781.

Fischl B, Rajendran N, Busa E, Augustinack J, Hinds O, Yeo BTT, Mohlberg H, Amunts K, Zilles K. 2008. Cortical Folding Patterns and Predicting Cytoarchitecture. Cereb Cortex. 18:1973–1980.

Fischl B, Sereno MI, Tootell RBH, Dale AM, others. 1999. High-resolution intersubject averaging and a coordinate system for the cortical surface. Hum Brain Mapp. 8:272–284.

Fischl B, Salat DH, Busa E, Albert M, Dieterich M, Haselgrove C, Van Der Kouwe A, Killiany R, Kennedy D, Klaveness S, Montillo A, Markris N, Rosen B, Dale AM. 2002. Whole brain segmentation: automated labeling of neuroanatomical structures in the human brain. Neuron. 33:341–355.

Fischl B, Van Der Kouwe A, Destrieux C, Halgren E, Ségonne F, Salat DH, Busa E, Seideman LJ, Goldstein J, Kennedy D, Caviness V, Markris N, Rosen B, Dale AM. 2004. Automatically parcellating the human cerebral cortex. Cereb Cortex. 14:11–22.

Fornito A, Zalesky A, Breakspear M. 2013. Graph analysis of the human connectome: Promise, progress, and pitfalls. Neuroimage. in press.

Fox MD, Corbetta M, Snyder AZ, Vincent JL, Raichle ME. 2006. Spontaneous neuronal activity distinguishes human dorsal and ventral attention systems. Proc Natl Acad Sci. 103:10046–10051.

Fox MD, Raichle ME. 2007. Spontaneous fluctuations in brain activity observed with functional magnetic resonance imaging. Nat Rev Neurosci. 8:700–711.

Fraley C, Raftery AE. 1998. How many clusters? Which clustering method? Answers via model-based cluster analysis. Comput J. 41:578–588.

Geman S. Geman D. 1984. Stochastic relaxation, Gibbs distributions, and the Bayesian restoration of images. IEEE Trans Pattern Anal Mach Intell. 6:721–741.

Geyer S. 2004. The Microstructural Border Between the Motor and the Cognitive Domain in the Human Cerebral Cortex, Advances in Anatomy Embryology and Cell Biology. Berlin, Heidelberg: Springer Berlin Heidelberg.

Geyer S, Ledberg A, Schleicher A, Kinomura S, Schormann T, Bürgel U, Klingberg T, Larsson J, Zilles K, Roland PE. 1996. Two different areas within the primary motor cortex of man.

Geyer S, Matelli M, Luppino G, Zilles K. 2000. Functional neuroanatomy of the primate isocortical motor system. Anat Embryol (Berl). 202:443–474.

Geyer S, Schleicher A, Schormann T, Mohlberg H, Bodegård A, Roland PE, Zilles K. 2001. Integration of microstructural and functional aspects of human somatosensory areas 3a, 3b, and 1 on the basis of a computerized brain atlas. Anat Embryol (Berl). 204:351–366.

Geyer S, Schleicher A, Zilles K. 1999. Areas 3a, 3b, and 1 of human primary somatosensory cortex: 1. Microstructural organization and interindividual variability. Neuroimage. 10:63–83.

Ghosh A, Rho Y, McIntosh a R, Kötter R, Jirsa VK. 2008. Noise during rest enables the exploration of the brain’s dynamic repertoire. PLoS Comput Biol. 4:e1000196.

Glahn DC, Winkler AM, Kochunov P, Almasy L, Duggirala R, Carless MA, Curran JC, Olvera RL, Laird AR, Smith SM, Beckmann CF, Fox PT, Blangero J. 2010. Genetic control over the resting brain. Proc Natl Acad Sci U S A. 107:1223–1228.

Glasser MF, Coalson TS, Robinson EC, Hacker CD, Harwell J, Yacoub E, Ugurbil K, Andersson J, Beckmann CF, Jenkinson M, Smith SM, Van Essen DC. 2016. A Multi-modal Parcellation of Human Cerebral Cortex. Nature.

Glasser MF, Sotiropoulos SN, Wilson JA, Coalson TS, Fischl B, Andersson JL, Xu J, Jbabdi S, Webster M, Polimeni JR, others. 2013. The minimal preprocessing pipelines for the Human Connectome Project. Neuroimage. 80:105–124.

Glasser MF, Van Essen DC. 2011. Mapping human cortical areas in vivo based on myelin content as revealed by T1-and T2-weighted MRI. J Neurosci. 31:11597–11616.

Gordon EM, Laumann TO, Adeyemo B, Huckins JF, Kelley WM, Petersen SE. 2016. Generation and Evaluation of a Cortical Area Parcellation from Resting-State Correlations. Cereb Cortex. 26:288–303.

Gordon EM, Laumann TO, Adeyemo B, Petersen SE. 2017. Individual Variability of the System-Level Organization of the Human Brain. Cereb Cortex. 27: 386–399

Greicius MD, Krasnow B, Reiss AL, Menon V. 2003. Functional connectivity in the resting brain: a network analysis of the default mode hypothesis. Proc Natl Acad Sci. 100:253–258.

Greve DN, Fischl B. 2009. Accurate and robust brain image alignment using boundary-based registration. Neuroimage. 48:63–72.

Griffanti L, Salimi-Khorshidi G, Beckmann CF, Auerbach EJ, Douaud G, Sexton CE, Zsoldos E, Ebmeier KP, Filippini N, Mackay CE, Moeller S, Xu J, Yacoub E, Baselli G, Ugurbil K, Miller KL, Smith SM. 2014. ICA-based artefact removal and accelerated fMRI acquisition for improved resting state network imaging. Neuroimage. 95:232–247.

Guye M, Bettus G, Bartolomei F, Cozzone PJ. 2010. Graph theoretical analysis of structural and functional connectivity MRI in normal and pathological brain networks. MAGMA.

Haber SN. 2003. The primate basal ganglia: parallel and integrative networks. J Chem Neuroanat. 26:317–330.

Hacker CD, Laumann TO, Szrama NP, Baldassarre A, Snyder AZ, Leuthardt EC, Corbetta M. 2013. Resting state network estimation in individual subjects. Neuroimage. 82:616–633.

Harrison SJ, Woolrich MW, Robinson EC, Glasser MF, Beckmann CF, Jenkinson M, Smith SM. 2015. Large-scale Probabilistic Functional Modes from resting state fMRI. Neuroimage. 109:217–231.

Hawrylycz M, Miller JA, Menon V, Feng D, Dolbeare T, Guillozet-Bongaarts AL, Jegga AG, Aronow BJ, Lee C-K, Bernard A, others. 2015. Canonical genetic signatures of the adult human brain. Nat Neurosci. 18:1832–1844.

He Y, Evans A. 2010. Graph theoretical modeling of brain connectivity. Curr Opin Neurol. 23:341–350.

Hill J, Inder T, Neil J, Dierker D, Harwell J, Van Essen D. 2010. Similar patterns of cortical expansion during human development and evolution. Proc Natl Acad Sci. 107:13135–13140.

Hinds O, Polimeni JR, Rajendran N, Balasubramanian M, Amunts K, Zilles K, Schwartz EL, Fischl B, Triantafyllou C. 2009. Locating the functional and anatomical boundaries of human primary visual cortex. Neuroimage. 46:915–922.

Hirose S, Watanabe T, Jimura K, Katsura M, Kunimatsu A, Abe O, Ohtomo K, Miyashita Y, Konishi S. 2012. Local Signal Time-Series during Rest Used for Areal Boundary Mapping in Individual Human Brains. PLoS One. 7:e36496.

Holmes AJ, Hollinshead MO, O’Keefe TM, Petrov VI, Fariello GR, Wald LL, Fischl B, Rosen BR, Mair RW, Roffman JL, Smoller JW, Buckner RL. 2015. Brain Genomics Superstruct Project initial data release with structural, functional, and behavioral measures. Sci data. 2:150031.

Honey CJ, Sporns O, Cammoun L, Gigandet X, Thiran JP, Meuli R, Hagmann P. 2009. Predicting human resting-state functional connectivity from structural connectivity. Proc Natl Acad Sci U S A. 106:2035–2040.

Honnorat N, Eavani H, Satterthwaite TD, Gur RE, Gur RC, Davatzikos C. 2015. GraSP: Geodesic Graph-based Segmentation with Shape Priors for the functional parcellation of the cortex. Neuroimage. 106:207–221.

Hubel DH, Wiesel TN. 1965. Receptive Fields and Functional Architecture in Two Nonstriate Visual Areas (18 and 19) of the Cat. J Neurophysiol. 28:229–289.

Iwamura Y. 1998. Hierarchical somatosensory processing. Curr Opin Neurobiol. 8:522–528.

Jiang L, Xu T, He Y, Hou X-H, Wang J, Cao X-Y, Wei G-X, Yang Z, He Y, Zuo X-N. 2015. Toward neurobiological characterization of functional homogeneity in the human cortex: regional variation, morphological association and functional covariance network organization. Brain Struct Funct. 220:2485–2507.

Jiang L, Zuo X-N. 2016. Regional homogeneity: a multimodal, multiscale neuroimaging marker of the human connectome. Neuroscientist. 22:486–505.

Jones DK, Symms MR, Cercignani M, Howard RJ. 2005. The effect of filter size on VBM analyses of DT-MRI data. Neuroimage. 26:546–554.

Jones EG. 1985. The Thalamus. Boston, MA: Springer US.

Jones EG. 1986. Connectivity of the primate sensory-motor cortex. In: Sensory-motor areas and aspects of cortical connectivity. Springer. p. 113–183.

Kaas J. 1987. The Organization Of Neocortex In Mammals: Implications For Theories Of Brain Function. Annu Rev Psychol. 38:129–151.

Krienen FM, Yeo BTT, Buckner RL. 2014. Reconfigurable task-dependent functional coupling modes cluster around a core functional architecture. Phil Trans R Soc B. 369:20130526.

Krienen FM, Yeo BTT, Ge T, Buckner RL, Sherwood CC. 2016. Transcriptional profiles of supragranular-enriched genes associate with corticocortical network architecture in the human brain. Proc Natl Acad Sci. 113:E469–E478.

Krubitzer L, Huffman KJ, Disbrow E, Recanzone G. 2004. Organization of area 3a in macaque monkeys: contributions to the cortical phenotype. J Comp Neurol. 471:97–111.

Kwong KK, Belliveau JW, Chesler DA, Goldberg IE, Weisskoff RM, Poncelet BP, Kennedy DN, Hoppel BE, Cohen MS, Turner R. 1992. Dynamic magnetic resonance imaging of human brain activity during primary sensory stimulation. Proc Natl Acad Sci. 89:5675–5679.

Lange T, Roth V, Braun ML, Buhmann JM. 2004. Stability-based validation of clustering solutions. Neural Comput. 16:1299–1323.

Laumann TO, Gordon EM, Adeyemo B, Snyder AZ, Joo SJ, Chen M-Y, Gilmore AW, McDermott KB, Nelson SM, Dosenbach NUF, others. 2015. Functional system and areal organization of a highly sampled individual human brain. Neuron. 87:657–670.

Lee MH, Hacker CD, Snyder AZ, Corbetta M, Zhang D, Leuthardt EC, Shimony JS. 2012. Clustering of resting state networks. PLoS One. 7:e40370.

Livingstone MS, Hubel DH. 1984. Anatomy and physiology of a color system in the primate visual cortex. J Neurosci. 4:309–356.

Long X, Goltz D, Margulies DS, Nierhaus T, Villringer A. 2014. Functional connectivity-based parcellation of the human sensorimotor cortex. Eur J Neurosci. 39:1332–1342.

Lotze M, Erb M, Flor H, Huelsmann E, Godde B, Grodd W. 2000. fMRI evaluation of somatotopic representation in human primary motor cortex. Neuroimage. 11: 473–481.

Lu J, Liu H, Zhang M, Wang D, Cao Y, Ma Q, Rong D, Wang X, Buckner RL, Li K. 2011. Focal pontine lesions provide evidence that intrinsic functional connectivity reflects polysynaptic anatomical pathways. J Neurosci. 31:15065–15071.

Malikovic A, Amunts K, Schleicher A, Mohlberg H, Eickhoff SB, Wilms M, Palomero-Gallagher N, Armstrong E, Zilles K. 2007. Cytoarchitectonic analysis of the human extrastriate cortex in the region of V5/MT+: a probabilistic, stereotaxic map of area hOc5. Cereb Cortex. 17:562–574.

Margulies DS, Kelly AMC, Uddin LQ, Biswal BB, Castellanos FX, Milham MP. 2007. Mapping the functional connectivity of anterior cingulate cortex. Neuroimage. 37:579–588.

Maunsell JHR, Van Essen DC. The connections of the middle temporal visual area (MT) and their relationship to a cortical hierarchy in the macaque monkey. J Neurosci 3: 2563, 1983.

Mennes M, Kelly C, Zuo X-N, Di Martino A, Biswal BB, Castellanos FX, Milham MP. 2010. Inter-individual differences in resting-state functional connectivity predict task-induced BOLD activity. Neuroimage. 50:1690–1701.

Meunier D, Achard S, Morcom A, Bullmore E. 2009. Age-related changes in modular organization of human brain functional networks. Neuroimage. 44:715–723.

Mueller S, Wang D, Fox MD, Pan R, Lu J, Li K, Sun W, Buckner RL, Liu H. 2015. Reliability correction for functional connectivity: Theory and implementation. Hum Brain Mapp. 36:4664–4680.

Mueller S, Wang D, Fox MD, Yeo BTT, Sepulcre J, Sabuncu MR, Shafee R, Lu J, Liu H. 2013. Individual variability in functional connectivity architecture of the human brain. Neuron. 77:586–595.

Nelson SM, Dosenbach NUF, Cohen AL, Wheeler ME, Schlaggar BL, Petersen SE. 2010. Role of the anterior insula in task-level control and focal attention. Brain Struct Funct. 214:669–680.

Nishitani N, Schürmann M, Amunts K, Hari R. 2005. Broca’s region: from action to language. Physiology. 20:60–69.

Nooner KB, Colcombe SJ, Tobe RH, Mennes M, Benedict MM, Moreno AL, Panek LJ, Brown S, Zavitz ST, Li Q, Sikka S, Gutman D, Bangaru S, Schlachter RT, Kamiel SM, Anwar AR, Hinz CM, Kaplan MS, Rachlin AB, Adelsberg S, Cheung B, Khanuja R, Yan C, Craddock CC, Calhoun V, Courtney W, King M, Wood D, Cox CL, Kelly AM, Di Martino A, Petkova E, Reiss PT, Duan N, Thomsen D, Biswal B, Coffey B, Hoptman MJ, Javitt DC, Pomara N, Sidtis JJ, Koplewicz HS, Castellanos FX, Leventhal BL, Milham MP. 2012. The NKI-Rockland Sample: A Model for Accelerating the Pace of Discovery Science in Psychiatry. Front Neurosci. 6:152.

Nowozin S, Lampert CH. 2010. Global Interactions in Random Field Models: A Potential Function Ensuring Connectedness. SIAM J Imaging Sci. 3:1048–1074.

Ogawa S, Tank DW, Menon R, Ellermann JM, Kim SG, Merkle H, Ugurbil K. 1992. Intrinsic signal changes accompanying sensory stimulation: functional brain mapping with magnetic resonance imaging. Proc Natl Acad Sci. 89:5951–5955.

Petersen SE, Fox PT, Posner MI, Mintun M, Raichle ME. 1988. Positron emission tomographic studies of the cortical anatomy of single-word processing. Nature. 331:585–589.

Peyron C, Petit J-M, Rampon C, Jouvet M, Luppi P-H. 1997. Forebrain afferents to the rat dorsal raphe nucleus demonstrated by retrograde and anterograde tracing methods. Neuroscience. 82:443–468.

Poldrack RA. 2006. Can cognitive processes be inferred from neuroimaging data? Trends Cogn Sci. 10:59–63.

Power JD, Cohen AL, Nelson SM, Wig GS, Barnes KA, Church JA, Vogel AC, Laumann TO, Miezin FM, Schlaggar BL, Petersen SE. 2011. Functional network organization of the human brain. Neuron. 72:665–678.

Preuss TM. 2004. What is it like to be a human. Cogn Neurosci. 3:5–22.

Raichle ME. 1987. Circulatory and metabolic correlates of brain function in normal humans. Compr Physiol.

Richiardi J, Altmann A, Milazzo A-C, Chang C, Chakravarty MM, Banaschewski T, Barker GJ, Bokde ALW, Bromberg U, Büchel C, others. 2015. Correlated gene expression supports synchronous activity in brain networks. Science (80−). 348:1241–1244.

Robinson EC, Jbabdi S, Glasser MF, Andersson J, Burgess GC, Harms MP, Smith SM, Van Essen DC, Jenkinson M. 2014. MSM: A new flexible framework for Multimodal Surface Matching. Neuroimage. 100:414–426.

Ryali S, Chen T, Supekar K, Menon V. 2013. A parcellation scheme based on von Mises-Fisher distributions and Markov random fields for segmenting brain regions using resting-state fMRI. Neuroimage. 65:83–96.

Salvador R, Suckling J, Coleman MR, Pickard JD, Menon D, Bullmore E. 2005. Neurophysiological architecture of functional magnetic resonance images of human brain. Cereb Cortex. 15:1332–1342.

Schaefer A, Margulies DS, Lohmann G, Gorgolewski KJ, Smallwood J, Kiebel SJ, Villringer A. 2014. Dynamic network participation of functional connectivity hubs assessed by resting-state fMRI. Front Hum Neurosci. 8:195.

Schmahmann JD, Pandya DN. 1997. Anatomic organization of the basilar pontine projections from prefrontal cortices in rhesus monkey. J Neurosci. 17:438–458.

Seeley WW, Menon V, Schatzberg AF, Keller J, Glover GH, Kenna H, Reiss AL, Greicius MD. 2007. Dissociable intrinsic connectivity networks for salience processing and executive control. J Neurosci. 27:2349–2356.

Sereno MI, Dale AM, Reppas JB, Kwong KK, Belliveau JW, Brady TJ, Rosen BR, Tootell RB. 1995. Borders of multiple visual areas in humans revealed by functional magnetic resonance imaging. Science. 268:889–893.

Shadlen MN, Newsome WT. 1998. The variable discharge of cortical neurons: implications for connectivity, computation, and information coding. The Journal of neuroscience, 18(10), pp.3870–3896.

Shen X, Tokoglu F, Papademetris X, Constable RT. 2013. Groupwise whole-brain parcellation from resting-state fMRI data for network node identification. Neuroimage. 82:403–415.

Smith SM, Beckmann CF, Andersson J, Auerbach EJ, Bijsterbosch J, Douaud G, Duff E, Feinberg DA, Griffanti L, Harms MP, Kelly M, Laumann T, Miller KL, Moeller S, Petersen S, Power J, Salimi-Khorshidi G, Snyder AZ, Vu AT, Woolrich MW, Xu J, Yacoub E, Ugurbil K, Van Essen DC, Glasser MF. 2013. Resting-state fMRI in the Human Connectome Project. Neuroimage. 80:144–168.

Smith SM, Fox PT, Miller KL, Glahn DC, Fox PM, Mackay CE, Filippini N, Watkins KE, Toro R, Laird AR, others. 2009. Correspondence of the brain’s functional architecture during activation and rest. Proc Natl Acad Sci. 106:13040–13045.

Smith SM, Nichols TE, Vidaurre D, Winkler AM, Behrens TE, Glasser MF, Ugurbil K, Barch DM, Van Essen DC, Miller KL. 2015. A positive-negative mode of population covariation links brain connectivity, demographics and behavior. Nat Neurosci, 18:1565–1567

Strick PL. 1985. How do the basal ganglia and cerebellum gain access to the cortical motor areas? Behav Brain Res. 18:107–123.

Strick PL, Dum RP, Fiez JA. 2009. Cerebellum and nonmotor function. Annu Rev Neurosci. 32:413–434.

Striem-Amit E, Ovadia-Caro S, Caramazza A, Margulies DS, Villringer A, Amedi A. 2015. Functional connectivity of visual cortex in the blind follows retinotopic organization principles. Brain. 138:1679–1695.

Supekar K, Menon V, Rubin D, Musen M, Greicius MD. 2008. Network analysis of intrinsic functional brain connectivity in Alzheimer’s disease. PLoS Comput Biol. 4:e1000100.

Tavor I, Jones OP, Mars RB, Smith SM, Behrens TE, Jbabdi S. 2016. Task-free MRI predicts individual differences in brain activity during task performance. Science (80−). 352:216–220.

Toga AW, Thompson PM, Mori S, Amunts K, Zilles K. 2006. Towards multimodal atlases of the human brain. Nat Rev Neurosci. 7:952–966.

Ungerleider LG, Desimone R. 1986. Cortical connections of visual area MT in the macaque. J Comp Neurol. 248:190–222.

van den Heuvel M, Mandl R, Hulshoff Pol H. 2008. Normalized Cut Group Clustering of Resting-State fMRI Data. PLoS One. 3:e2001.

Van Essen DC, Zeki SM. The topographic organization of rhesus monkey prestriate cortex. J Physiol 277: 193–226, 1978.

Van Essen DC, Dierker DL. 2007. Surface-based and probabilistic atlases of primate cerebral cortex. Neuron. 56:209–225.

Van Essen DC, Glasser MF. 2014. In vivo architectonics: a cortico-centric perspective. Neuroimage. 93:157–164.

Van Essen DC, Glasser MF, Dierker DL, Harwell J, Coalson T. 2012a. Parcellations and hemispheric asymmetries of human cerebral cortex analyzed on surface-based atlases. Cereb Cortex. 22:2241–2262.

Van Essen DC, Ugurbil K, Auerbach E, Barch D, Behrens TEJ, Bucholz R, Chang A, Chen L, Corbetta M, Curtiss SW, Della Penna S, Feinberg D, Glasser MF, Harel N, Heath AC, Larson-Prior L, Marcus D, Michalareas G, Moeller S, Oostenveld R, Petersen SE, Prior F, Schlaggar BL, Smith SM, Snyder AZ, Xu J, Yacoub E. 2012b. The Human Connectome Project: A data acquisition perspective. Neuroimage. 62:2222–2231.

Varoquaux G, Gramfort A, Pedregosa F, Michel V, Thirion B. 2011. Multi-subject Dictionary Learning to Segment an Atlas of Brain Spontaneous Activity. In: Information Processing in Medical Imaging. p. 562–573.

Vincent JL, Kahn I, Snyder AZ, Raichle ME, Buckner RL. 2008. Evidence for a frontoparietal control system revealed by intrinsic functional connectivity. J Neurophysiol. 100:3328–3342.

Vincent JL, Patel GH, Fox MD, Snyder a Z, Baker JT, Van Essen DC, Zempel JM, Snyder LH, Corbetta M, Raichle ME. 2007. Intrinsic functional architecture in the anaesthetized monkey brain. Nature. 447:83–86.

Vincent JL, Snyder AZ, Fox MD, Shannon BJ, Andrews JR, Raichle ME, Buckner RL. 2006. Coherent spontaneous activity identifies a hippocampal-parietal memory network. J Neurophysiol. 96:3517–3531.

Vogt B. 2009. Cingulate neurobiology and disease. Oxford University Press.

Vogt C, Vogt O. 1919. Allgemeine ergebnisse unserer hirnforschung. JA Barth.

von Economo CF, Koskinas GN. 1925. Die cytoarchitektonik der hirnrinde des erwachsenen menschen. J. Springer.

Wandell BA, Dumoulin SO, Brewer AA. 2007. Visual field maps in human cortex. Neuron. 56:366–383.

Wang J, Fan L, Wang Y, Xu W, Jiang T, Fox PT, Eickhoff SB, Yu C, Jiang T. 2015a. Determination of the posterior boundary of Wernicke’s area based on multimodal connectivity profiles. Hum Brain Mapp. 36:1908–1924.

Wang J, Yang Y, Fan L, Xu J, Li C, Liu Y, Fox PT, Eickhoff SB, Yu C, Jiang T. 2015b. Convergent functional architecture of the superior parietal lobule unraveled with multimodal neuroimaging approaches. Hum Brain Mapp. 36:238–257.

Wang J, Zuo X, He Y. 2010. Graph-based network analysis of resting-state functional MRI. Front Syst Neurosci. 4:16.

Wang Q, Sporns O, Burkhalter A. 2012. Network analysis of corticocortical connections reveals ventral and dorsal processing streams in mouse visual cortex. J Neurosci. 32:4386–4399.

Wig GS, Laumann TO, Cohen AL, Power JD, Nelson SM, Glasser MF, Miezin FM, Snyder AZ, Schlaggar BL, Petersen SE. 2014a. Parcellating an individual subject’s cortical and subcortical brain structures using snowball sampling of resting-state correlations. Cereb Cortex. 24:2036–2054.

Wig GS, Laumann TO, Petersen SE. 2014b. An approach for parcellating human cortical areas using resting-state correlations. Neuroimage. 93:276–291.

Xu T, Opitz A, Craddock RC, Zuo X-N, Milham M. 2016. Assessing variations in areal organization for the intrinsic brain: From fingerprints to reliability. Cereb Cortex. 26:4192–4211.

Xu T, Yang Z, Jiang L, Xing X-X, Zuo X-N. 2015. A connectome computation system for discovery science of brain. Sci Bull. 60:86–95.

Yang Z, Chang C, Xu T, Jiang L, Handwerker DA, Castellanos FX, Milham MP, Bandettini PA, Zuo X-N. 2014. Connectivity trajectory across lifespan differentiates the precuneus from the default network. Neuroimage. 89:45–56.

Yang Z, Zuo X-N, McMahon KL, Craddock RC, Kelly C, de Zubicaray GI, Hickie I, Bandettini PA, Castellanos FX, Milham MP, others. 2016. Genetic and Environmental Contributions to Functional Connectivity Architecture of the Human Brain. Cereb Cortex. 26:2341–2352.

Yeo BTT, Eickhoff SB. 2016. A modern map of the human cerebral cortex. Nature. 536:152–154

Yeo BTT, Krienen FM, Chee MWL, Buckner RL. 2014. Estimates of segregation and overlap of functional connectivity networks in the human cerebral cortex. Neuroimage. 88:212–227.

Yeo BTT, Krienen FM, Eickhoff SB, Yaakub SN, Fox PT, Buckner RL, Asplund CL, Chee MWL. 2015a. Functional Specialization and Flexibility in Human Association Cortex. Cereb Cortex. 25:3654–3672.

Yeo BTT, Krienen FM, Sepulcre J, Sabuncu MR, Lashkari D, Hollinshead M, Roffman JL, Smoller JW, Zöllei L, Polimeni JR, Fischl B, Liu H, Buckner RL. 2011. The organization of the human cerebral cortex estimated by intrinsic functional connectivity. J Neurophysiol. 106:1125–1165.

Yeo BTT, Sabuncu MR, Vercauteren T, Ayache N, Fischl B, Golland P. 2010. Spherical demons: fast diffeomorphic landmark-free surface registration. IEEE Trans Med Imaging. 29:650–668.

Yeo BTT, Tandi J, Chee MWL. 2015b. Functional connectivity during rested wakefulness predicts vulnerability to sleep deprivation. Neuroimage. 111:147–158.

Zhang Y, Brady M, Smith S. 2001. Segmentation of brain MR images through a hidden Markov random field model and the expectation-maximization algorithm. IEEE Trans Med Imaging. 20: 45–57.

Zalesky A, Fornito A, Bullmore ET. 2010a. Network-based statistic: Identifying differences in brain networks. Neuroimage. 53:1197–1207.

Zalesky A, Fornito A, Cocchi L, Gollo LL, Breakspear M. 2014. Time-resolved resting-state brain networks. Proc Natl Acad Sci. 111:10341–10346.

Zalesky A, Fornito A, Harding IH, Cocchi L, Yücel M, Pantelis C, Bullmore ET. 2010b. Whole-brain anatomical networks: does the choice of nodes matter? Neuroimage. 50:970–983.

Zeki SM. Representation of central visual fields in prestriate cortex of monkey. Brain Res 14: 271–291, 1969.

Zuo X-N, Ehmke R, Mennes M, Imperati D, Castellanos FX, Sporns O, Milham MP. 2012. Network Centrality in the Human Functional Connectome. Cereb Cortex. 22:1862–1875.

Zuo X-N, He Y, Betzel RF, Colombe S, Sporns O, Milham MP. 2017. Human connectomics across the lifepsan. Trends in Cognitive Sciences. 21: 32–45.

Zuo X-N, Kelly C, Adelstein JS, Klein DF, Castellanos FX, Milham MP. 2010. Reliable intrinsic connectivity networks: test-retest evaluation using ICA and dual regression approach. Neuroimage. 49:2163–2177.

Zuo X-N, Xing X-X. 2014. Test-retest reliabilities of resting-state FMRI measurements in human brain functional connectomics: a systems neuroscience perspective. Neurosci Biobehav Rev. 45:100–118.

Zuo X-N, Xu T, Jiang L, Yang Z, Cao X-Y, He Y, Zang Y-F, Castellanos FX, Milham MP. 2013. Toward reliable characterization of functional homogeneity in the human brain: preprocessing, scan duration, imaging resolution and computational space. Neuroimage. 65:374–386.

